# Data-driven traction force microscopy in 3D collagen hydrogels

**DOI:** 10.1101/2025.11.17.688772

**Authors:** Jorge Barrasa-Fano, Laurens Kimps, Alejandro Apolinar-Fernández, Débora Linhares, Iain Muntz, Nele Vaes, Apeksha Shapeti, Elías Núñez-Ortega, Ruth Cardinaels, Gijsje H. Koenderink, José Antonio Sanz-Herrera, Hans Van Oosterwyck

## Abstract

Three-dimensional traction force microscopy (3DTFM) in fibrillar hydrogels typically models the extracellular matrix as a continuum. We present data-driven 3DTFM (DD-TFM), which reconstructs fiber networks from confocal images and models them as discrete fiber networks mechanically calibrated by shear rheology. We validate DD-TFM in silico and apply it to endothelial cells, quantifying cell-generated forces and matrix stresses at the fiber level. Compared with continuum TFM DD-TFM reveals network-guided matrix deformations, shifting 3DTFM toward fiber-resolved force calculation.

## Main

Cells live in and interact with the extracellular matrix (ECM), a complex scaffold of biomolecules that varies in composition and organization across tissues^1^. The ECM’s mechanical properties influence key physiological and pathological processes such as cell migration^2^, morphogenesis^3^ and cancer progression^4^. Cells also actively probe and remodel the ECM by generating forces and transmitting them through cell-ECM adhesions, enabling mechanosensing and mechanotransduction^5^.

Fibrous hydrogels like collagen type I and fibrin are widely used to mimic the 3D ECM in vitro, given their physiological relevance, cytocompatibility, and nonlinear elastic behavior^6,7^. They support integrin-based adhesion and matrix remodeling through matrix degrading enzymes^8^, making them suitable for applications ranging from angiogenesis^9^ and stem cell differentiation^10^ to cancer invasion and metastasis^11^. Over the past 16 years^12^, the mechanobiology community has developed methods to quantify cellular forces in 3D hydrogels. Traction force microscopy (TFM)^13^ was initially established for 2D linear elastic substrates^14^ and later extended to 3D using the finite element method (FEM) in linear polyethylene glycol (PEG) hydrogels^12^. However, applying FEM to nonlinear fibrous matrices like collagen is technically challenging. Many have measured matrix deformations alone^9,15–19^ or assumed simplified material behaviors such as linear elastic^20–22^ or Neo-Hookean material models^23^. Pioneered by the seminal work of Steinwachs et al.^24^, the most advanced nonlinear 3D TFM works have developed FEM-implemented constitutive models that incorporate buckling, straightening and stretching regimes of the bulk mechanical behavior of the fibrous network^24–28^. These studies consider the ECM as a continuum with homogeneous mechanical properties often using a bulk mechanical characterization of hydrogel samples. In continuum mechanics, the representative volume element (RVE) used to theoretically derive constitutive models for materials with an internal microstructure serves as the minimal volume over which mechanical properties can be homogenized. However, this essential assumption inherently limits the resolution at which mechanical heterogeneities can be modeled and imposes smooth displacement fields with spatial decay governed by the homogenized material response. In fibrous hydrogels, composed of discrete fibers, the existence of a meaningful RVE is not always guaranteed, particularly at the (sub)cellular scale at which forces are applied and sensed^29^. As a result, continuum models may oversimplify or misrepresent the true mechanical microenvironment that cells sense and create. Recently, Ucla et al. performed 3DTFM at fiber level but relied on simplified photopolymerized synthetic fiber arrays^30^. To overcome these limitations, we propose a data-driven, discrete fiber-based approach to 3DTFM (DD-TFM). Our method obtains fibrous matrix architecture directly from confocal microscopy images and models the ECM as a network of interconnected fibers (**Figure 1**a). By replacing continuum homogenization with image-derived discrete fiber networks, this method resolves how cells deform their fibrous surroundings at sub-RVE length scales (**Figure 1**f-i), representing a paradigm shift for 3DTFM.

**Figure 1:**
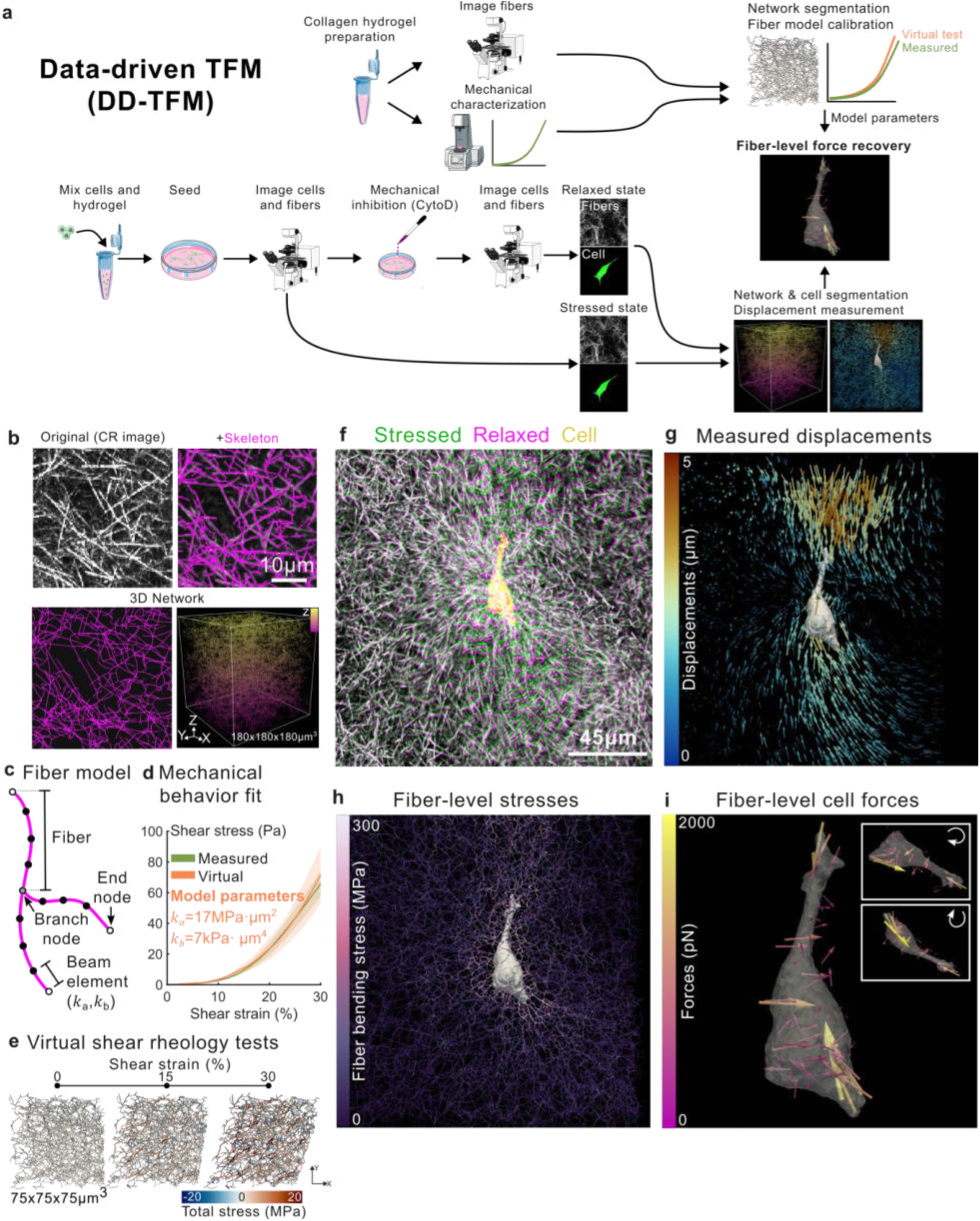
The data driven TFM (DD-TFM) pipeline captures cell forces and matrix stresses at the fiber level. **a:** Basic schematic of DD-TFM. Top flow: collagen samples are prepared to undergo microscopy imaging (**b**-top left: maximum projection of 35 slices of a confocal reflectance (CR) image stack) and shear rheology tests (**d**-green line: n = 3). Microscopy images are segmented and skeletonized (**b**-top right) to build a graph-based representation of the 3D network architecture (**b**-bottom). Fibers in this model are represented as beam elements with axial stiffness (k_a_) and bending stiffness (k_b_) (**c**). These parameters are adjusted by performing virtual shear rheology on the segmented matrices and fitting their values to the measured curve (**d**-orange line: n = 4 sub-volumes and **e**: 3 representative steps of a virtual shear rheology test on a sub-volume extracted from the segmented matrix at applied shear strains γ=0, 15 and 30%. Here γ denotes the imposed simple-shear deformation. Fiber are color-coded by the model-derived element-wise total fiber stress which combines axial and bending stress contributions; colorbar limits adjusted to ease visualization. Bottom flow: TFM samples are prepared by mixing cells with a collagen hydrogel and 3D imaging them before (stressed state) and after (relaxed state) mechanical inhibition with Cytochalasin D (**f:** pseudocolor overlay of a maximum intensity projection of the cell (CellTracker), stressed, and relaxed state images (CR) in yellow, green and magenta, respectively; white indicates perfect overlap between the two states). Fiber images are registered to measure 3D matrix displacements (**g**: render of the displacement field colored by vector magnitude around the segmented cell) and segmented to build the network architecture model. Through an inverse method (HPBIM) displacements are translated into fiber level fiber stresses (**h**: color-coded fiber bending stresses at a cross-section of the matrix; colormap adjusted to ease visualization) and cell forces (**i**: force field colored by vector magnitude around the segmented cell and two additional rotated views (insets)).

When collagen hydrogels are used, we and others have measured matrix displacements by directly imaging matrix fibers (label-free either with second harmonic generation, or confocal reflectance (CR), or by fluorescently labeling the fibers) without the need for fluorescent beads^15,24,31^. DD-TFM uses these fiber images not only for measuring matrix deformations, but also to extract the underlying matrix architecture. While here we used CR as an illustrative example for collagen I hydrogels, our methodology is not limited to this imaging modality and can be applied using other techniques. We first segmented different ROIs of real CR images to build graph-based discrete fiber networks (**Figure 1**a,b; **Network segmentation from microscopy images**) in which fibers are modeled as chains of beams (**Figure 1**c) with axial stiffness, *k*_*a*_, and bending stiffness, *k*_*b*_. Then, we performed shear rheology tests on collagen hydrogel samples (**Figure 1**a,d-green curve; **Bulk shear rheology**). To guarantee that our model reproduces the macroscopic mechanics of our collagen hydrogels, we searched for the values *k*_*a*_ and *k*_*b*_ that best fit these experimental curves by performing virtual shear rheology tests on the segmented networks (**Figure 1**e, d-orange curve), obtaining *k*_*a*_=17 MPa·µm^2^ and *k*_*b*_=7 kPa· µm^4^, in line with the order of magnitude reported via AFM measurements^32,33^ (**Virtual shear rheology and parameter calibration**).

To identify the most robust force-recovery strategy for DD-TFM, we simulated a contracting cell in a real collagen network, generated ground-truth displacements and forces, and corrupted the displacement field with increasing levels of noise (**In silico ground truth simulations, Supplementary Videos 1**,**2**). A hybrid physics-based inverse method (*HPBIM*)^34,35^ and a classical Tikhonov-regularized (*Classic*) method, previously established for continuum 3D TFM, were extended to the treatment of discrete fiber networks and compared in terms of force recovery (**Recovery of cell forces**). *HPBIM* outperformed *Classic* across all metrics, maintaining force errors below 40% for realistic noise levels and preserving high spatial force correlations, whereas *Classic* underestimated force magnitudes and introduced non-physical forces away from the cell (**Extended Data Figure 3**).

To fully explore the potential of DD-TFM with *HPBIM*, we recovered the forces exerted by control (CT) or 15µM blebbistatin (Blebb) treated human umbilical vein endothelial cells (HUVECs) embedded in a collagen matrix (15 and 20 cells per condition, respectively; **Figure 2**a-top; **Cell culture and TFM sample preparation**). Cells presented an overall contractile behavior, with exerted forces typically pointing inwards (**Figure 2**a-bottom and positive total contractility values in **Figure 2**b-right). CT cells consistently induced larger fiber displacements (**Figure 2**a-middle) and *HPBIM* retrieved larger CT forces (**Figure 2**a-bottom, **Extended Data Figure 4**a-d), strain energy (internal work developed by cells to deform the fibers), total force and contractility compared to Blebb cells (**Figure 2**b). CT cells also induced higher total fiber stresses that propagated over longer distances away from the cell (**Figure 2**c).

**Figure 2:**
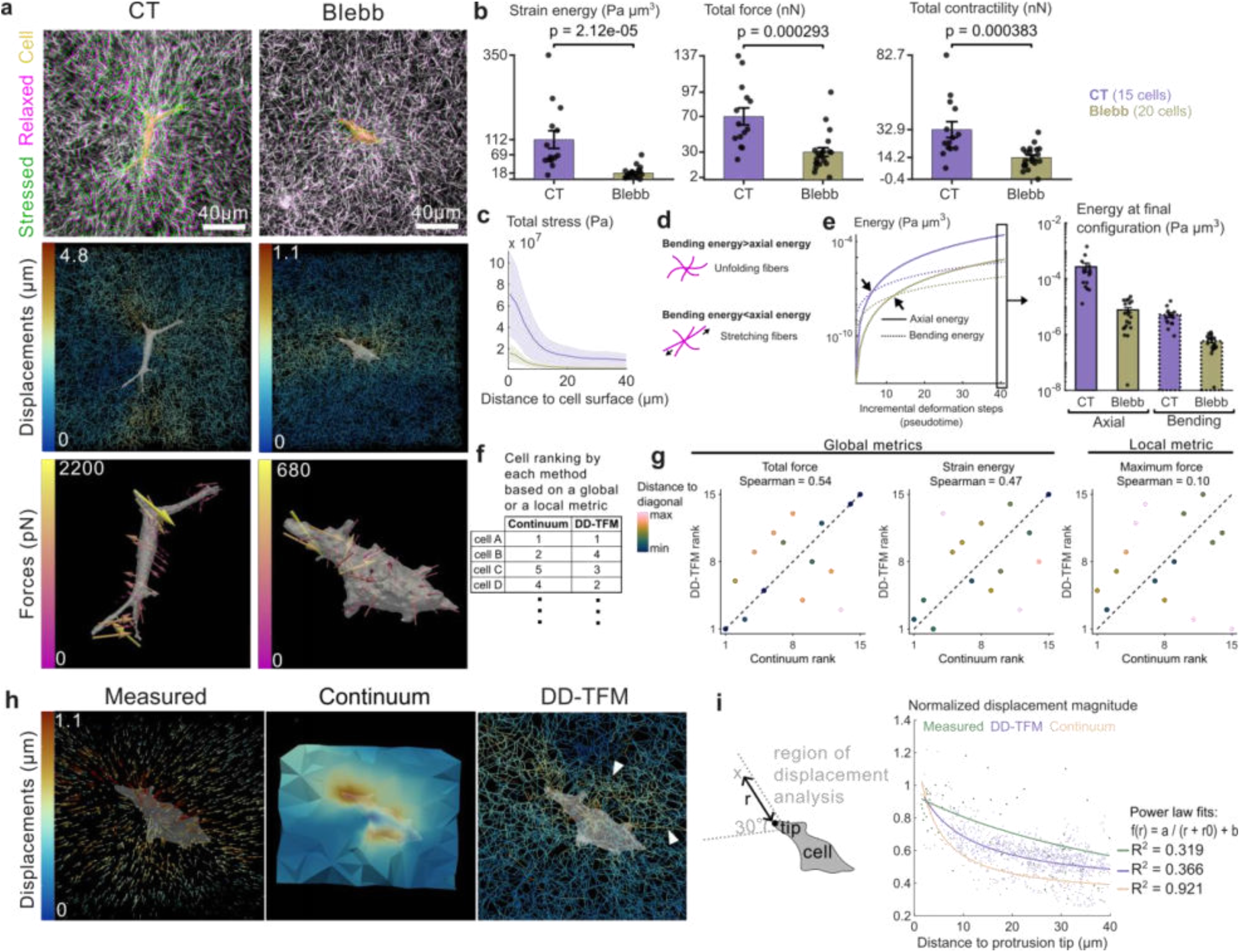
Fiber-level endothelial forces and network-guided matrix deformation with DD-TFM. **a:** (top row) Maximum intensity projections of stressed, relaxed CR images and the cell (labelled with CellTracker) image (green, magenta and yellow, respectively) for control (CT) and Blebbistatin-treated HUVECs (Blebb). (middle row) Color-coded fiber HPBIM displacement magnitude recovered at a cross-section of the matrix around segmented cells. (bottom row) Force field computed with HPBIM colored by vector magnitude around the segmented cells. **b**: Comparison of CT and Blebb cells across force metrics strain energy (left), total force (middle), and total contractility (right). Bars show mean ± SEM with individual cells overlaid as scatter points (15 cells for CT and 20 cells for Blebb). Statistical analysis was done using a two-sided Wilcoxon rank-sum test. **c**: Total fiber stress decay as a function of distance to the cell surface for CT and Blebb cells. Lines and shading indicate mean ± SD. **d:** Schematic representation of axial and bending deformation regimes in the discrete fiber model. **e:** Left, evolution of median axial and bending energy within 5 µm of the cell surface across incremental HPBIM deformation steps (pseudotime). Arrows indicate the transition from an initially bending-dominated regime to a stretching-dominated regime. Right, final axial and bending energy contributions, showing that both CT and Blebb cells reached a stretching-dominated state, with higher total matrix energy in CT cells. **f**: based on global or local force metrics, continuum and DD-TFM rank cells of the dataset in different order. **g:** Cell ranking comparison between continuum TFM and DD-TFM for total force, strain energy and maximum force in the subset of cells analyzed with both methods. Rank=1 corresponds to the highest metric value. Spearman’s rank correlation was used to assess agreement in cell ordering between methods. **h:** Representative comparison of displacement fields (color-coded by displacement magnitude) around a cell (in white) for measured (left), continuum TFM (middle) and DD-TFM (right) displacements. Measured displacements are displayed as vector arrows plotted on the image grid, continuum displacements are displayed on a cross-section of the FE mesh, and DD-TFM displacements are shown on a cross-section of the discrete fiber model. White arrows indicate fibers that, while being at a certain distance from the cell, undergo higher displacements than their neighboring fibers, which highlights force transmission along connected fiber paths. **i:** Left, schematic of the conical region used to analyze displacement decay away from a cell protrusion tip. r is the Euclidean distance of a point in the conical region to the protrusion tip. Right, normalized displacement magnitude as a function of Euclidean distance from the protrusion tip. R^2^ values for measured (green), DD-TFM (purple) and continuum (orange) displacements are indicated; RMSE values are 0.1812, 0.0992 and 0.0597, respectively.

Previous discrete fiber-network simulations predicted that contractile cells deform fibrous matrices through two mechanical regimes: an initial bending-dominated response followed by a stretching-dominated regime beyond a threshold deformation^36^. We therefore asked whether DD-TFM could capture this transition experimentally by decomposing the recovered matrix strain energy into axial and bending contributions (**Fiber stresses and energies in DD-TFM**). Across incremental deformation steps, all cells displayed two mechanical regimes: bending energy initially dominated over axial energy, whereas axial energy dominated beyond a threshold deformation that was reached earlier by CT cells (**Figure 2**d, e-left). Both CT and Blebb cells reached a final stretching-dominated regime, although CT cells stored higher total energy in the matrix (**Figure 2**e-right). These results illustrate how DD-TFM can resolve not only how strongly cells pull on the matrix, but also how DD-TFM can be used to validate theoretical predictions on mechanisms that govern a cell’s mechanical interactions with a fibrous matrix.

We next compared DD-TFM with a conventional continuum TFM approach for readouts that can be obtained with both methods (**Comparison of DD-TFM with continuum**). In a subset of 15 cells, we ranked cells according to global force readouts, including total force and strain energy, and a local readout, maximum force (**Figure 2**f, g). DD-TFM and continuum rankings were moderately correlated for global metrics (Spearman ~0.5) but weakly correlated for the local metric (Spearman = 0.1) (**Figure 2**g). Thus, continuum TFM can capture some global trends in cell-matrix mechanical interactions but may lead to different conclusions when local force values are considered, precisely where fiber-scale architecture is expected to matter most.

We then noticed that, overall, matrix stresses decayed faster in the continuum model than in DD-TFM (**Extended Data Figure 4**f). Although both continuum and DD-TFM used the same experimentally measured displacement field and *HPBIM* formulation, they differed in how the collagen matrix was represented: as a homogenized continuum or as an image-derived discrete fiber network. Each representation imposes different mechanical constraints on the recovered solution, leading to model-consistent displacement fields that can differ from the measured displacement field in distinct ways. Therefore, we compared the measured displacement field with the displacement fields obtained by continuum TFM and DD-TFM for a representative cell (**Figure 2**h). We fitted displacement magnitudes as a function of Euclidean distance r from a protrusion tip using a fixed inverse-distance model (**Figure 2**i-left). The continuum displacement field was well described by this smooth distance-dependent decay (R^2^=0.921), whereas both the measured displacements and DD-TFM displacements showed much weaker agreement (R^2^~0.3) (**Figure 2**i-right). This difference persisted when fitting a free-exponent power law (**Extended Data Figure 4**h). These results indicate that continuum TFM represents cell-induced matrix displacements as an effectively homogeneous, distance-dependent decay from the protrusion, whereas DD-TFM is not constrained in this way. Instead, nodes at similar distances from the cell can undergo different displacements depending on whether they are mechanically connected to the fibers being pulled (white arrows in **Figure 2**h). DD-TFM therefore captures a connectivity-guided displacement field, in which force transmission follows the local fiber architecture rather than Euclidean distance alone, more closely reflecting the heterogeneous displacement patterns measured in collagen matrices. By preserving these fiber-scale force-transmission pathways, DD-TFM may be particularly useful for studying long-range ECM-mediated mechanical communication between neighboring cells^37^.

This data-driven TFM framework bridges structural and mechanical information at the single-fiber level, providing a physically grounded view of how cells sense and mechanically interact with their 3D microenvironment. By resolving forces, stresses and fiber-level deformation modes within real collagen networks, DD-TFM extends beyond continuum approximations and enables quantification of how local matrix architecture governs cell-matrix mechanics. Such information is essential at (sub)micron length scales, where matrix architecture, focal adhesion dynamics and cytoskeletal remodeling interact below the length scale at which continuum descriptions are valid. By recovering cell-generated forces directly within image-derived fiber networks, DD-TFM shifts 3D TFM from continuum-scale force calculation toward fiber-level measurements of cell-ECM mechanotransduction.

## Methods

### Cell culture and TFM sample preparation

Human umbilical vein endothelial cells (HUVECs; Angio-Proteomie, Boston, MA) were cultured in complete endothelial growth medium (EGM-2, CC-3162, Lonza, Basel, Switzerland) supplemented with 100U/ml penicillin and 100μg/ml streptomycin on 100µg/mL collagen type I coated P60 Petri dishes. At passage 4, complete growth medium was replaced by 0.0125% (vol/vol) CellTracker Green CMFDA Dye (C7025, Invitrogen, Thermo Fisher Scientific, US) in supplemented growth medium and kept in the incubator for 20 minutes. During this time the collagen hydrogel mixture (see **Hydrogel preparation**) was prepared after which the cell suspension was pipetted into it for a resulting cell density of 10^4^ cells/mL. 100µL of the collagen mixture with cells was pipetted into wells of a glass bottom 96 well plate (#1.5H glass, IBL, Gerasdorf, Austria) while still on ice and immediately moved into the incubator at 37°C with 95% relative humidity, 20% O_2_ and 5% CO_2_ for 1h to allow for collagen polymerization. Finally, 250 µL of cell culture medium was added onto the polymerized gel and kept in the incubator at 37°C overnight. Treated cells were exposed overnight to blebbistatin at 15 µM (stock at 10mM dissolved in DMSO; B0560, Merck, Darmstadt, Germany) in supplemented growth medium.

### Hydrogel preparation

1.5 mg/mL type I collagen hydrogels were prepared based on previous work^38^. Collagen mixtures were prepared using 4°C reagents in Eppendorf tubes on ice to inhibit premature polymerization. Bovine hide derived type I atelocollagen (FibriCol®, 5133, Advanced Biomatrix, Carlsbad, CA, USA) was brought to a pH of 7.4 by addition of 1% (vol/vol) 0.1 M NaOH. Next, 10X PBS (Thermo Fisher Scientific, Waltham, MA, USA) was used to regulate osmolarity and salt concentrations. Finally, sterile milliQ water and cell suspension were added to reach a final collagen concentration of 1.5 mg/mL. For rheology experiments, the cell suspension was replaced by pure growth medium without cells. The collagen mixture was removed from ice and polymerized for one hour at 37°C on the rheometer plate (for shear rheology measurements) or in the incubator (for TFM experiments). Final volumes of 1000µL were prepared for both rheology (~150µL are needed for each rheometer measurement) and TFM measurements (>100µL/well are needed). For TFM measurements, fresh growth medium was added after one hour of polymerization.

### TFM data acquisition

Images were acquired on a confocal microscope (TCS SP8 DMi8, Leica Microsystems, Wetzlar, Germany) equipped with a 0.95 NA Fluotar VISIR 25x water immersion objective (Leica). Phase corrected bidirectional scanning at 2.5X zoom and a 55.9 µm pinhole (1 AU) created images with 512 x 512 resolution and voxel sizes of 0.36 x 0.36 x 1.0 µm^3^. Confocal imaging relied on simultaneous excitation of the cell channel at 488 nm and the fiber channel at 638 nm using a diode laser, with simultaneous detection of fluorescence emission at 495-555 nm (cell channel) and confocal reflectance signal at 631-644 nm (fiber channel) using a PMT and hybrid detector respectively. While it has been reported that confocal reflectance does not detect fibers oriented >50° relative to the imaging plane^39^, our methodology is not limited to this imaging modality and can be applied using other techniques. For TFM, the stressed state image was acquired first by marking the positions and then imaging the volumes of interest. A one-hour treatment with Cytochalasin D (4 µM, dissolved in DMSO; Sigma Aldrich, St. Louis, U.S.) was used to mechanically relax the cells. The same positions were imaged again to obtain the relaxed state. All imaging was performed at 37 °C in a humidified 5% CO2 environment. Our TFM dataset contains 35 cells (15 CT and 20 Blebb).

### Bulk shear rheology

Rheology tests were performed with a stress-controlled rotational rheometer (Physica MCR 702, Anton Paar, Graz, Austria) using a smooth stainless steel CP25-2 cone-plate geometry (25 mm diameter and 2° cone angle). 150 µL of cold collagen solution was pipetted onto the bottom plate preheated to 37 °C and the cone was immediately lowered. Mineral oil was added around the sample to prevent evaporation. A small-amplitude oscillatory shear test (1% strain, 1Hz) was performed for 1.5h (1 measured point every 60s) to measure the evolution of the storage and loss moduli with time. Then, a stress-controlled frequency sweep test (1% strain, frequency increasing from 0.1Hz to 10Hz) was performed. Finally, a stress-controlled rotation test (stresses logarithmically increasing from 0.01Pa to 100Pa, 10 measured strain points per decade, with 5s between each point) was performed for 3.5 minutes to obtain the stress-strain curve. Data was averaged over 3 independent replicates.

### Discrete fiber model

To virtually represent collagen networks we used the framework of the discrete fiber model we described previously^40^. Here, *fibers* are discretized by *nodes* (see **Figure 1**c). Nodes at the fiber ends (connectivity z=1) are called *end nodes*. Nodes of a fiber are connected through *elements* modeled as 3D Euler-Bernoulli beams with six degrees of freedom per node (three translations and three rotations) within an updated Lagrangian framework^40^. *Branch nodes* are nodes where more than 2 elements meet (connectivity z>2). The resultant multiscale model enables the analysis of micromechanical interactions between fibers and their contribution to the emergent macroscopic mechanical response of the matrix^41^. The mechanical response of individual fibers in this discrete network model is governed by axial stiffness, *k*_*a*_, bending stiffness, *k*_*b*_, and torsional stiffness, *k*_*t*_. These parameters can be defined as:

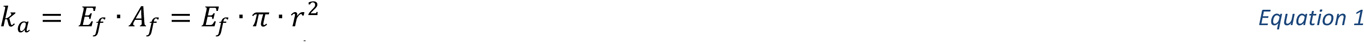

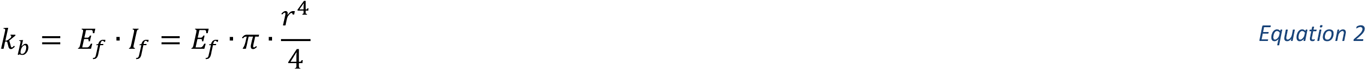

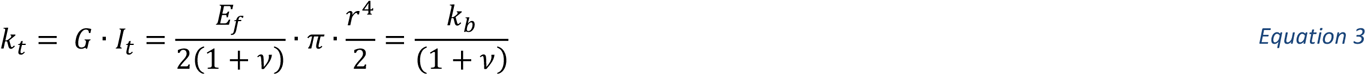

Where *E*_*f*_ is the fiber Young’s modulus, *A*_*f*_ is the cross-sectional area of the fibers, *I*_*f*_ is the second moment of inertia of the fiber cross-section, *I*_*t*_ is the polar moment of inertia, *G* is the shear modulus, *v* is the Poisson’s ratio and *r* is the fiber radius. The values used for these parameters are described in **Virtual shear rheology and parameter calibration**.

### Network segmentation from microscopy images

Three-dimensional fiber networks were reconstructed from volumetric microscopy images using an automated image-processing workflow implemented in MATLAB (**Extended Data Figure 1**a). First, the raw image stack was normalized to the intensity range [0,1] and denoised using a three-dimensional coherence-enhancing anisotropic diffusion filter. The filter was based on the Hybrid Diffusion with Continuous Switch (HDCS) formulation, which combines coherence-enhancing and edge-enhancing diffusion to smooth image noise while preserving elongated tubular structures and fiber continuity^42^. We used its MATLAB implementation `Image Edge Enhancing Coherence Filter Toolbox`^43^ with a total diffusion time of 1.5, a time step of 0.15, Gaussian pre-smoothing parameter σ = 0.1 voxel, structure tensor smoothing parameter ρ = 1 voxel, and contrast parameters λ_e_ = 0.02, λ_c_ = 0.02 and λ_h_ = 0.5. The filtered image was subsequently segmented into a binary fiber mask using adaptive local thresholding using MATLAB’S function `imbinarize` with a sensitivity parameter of 0.35. To suppress low-intensity background structures, only voxels with intensities above the 90th percentile of the filtered image were retained. Small isolated connected components containing fewer than 30 voxels were removed using morphological size filtering.

The binary mask was skeletonized using three-dimensional thinning (MATLAB `bwskel`) with a minimum branch length of 5 voxels. To compensate for incomplete skeletonization near fiber termini, skeleton endpoints were iteratively extended along their local orientation. For each endpoint, the local fiber direction was estimated from neighboring skeleton voxels within a radius of five voxels, and the endpoint was propagated up to ten voxels while remaining inside the segmented fiber mask. The resulting skeleton was converted into a graph representation using a topology-preserving skeleton-to-graph algorithm derived from the `Skel2Graph3D` framework^44^. Skeleton voxels with more than two neighbors were identified as branch nodes, voxels with a single neighbor as endpoints, and voxels with two neighbors as fiber segments connecting nodes. Connected voxel paths between neighboring nodes were extracted as graph links. Each extracted fiber path was transformed into physical coordinates using the image voxel size and resampled to obtain approximately uniform element lengths. Fiber centerlines were smoothed by fitting constrained second-order polynomials to the original voxel coordinates while preserving the positions of the fiber endpoints. The resulting curves were discretized at a target minimum element length, generating a graph whose nodes corresponded to spatial coordinates and whose edges represented fiber elements. Duplicate nodes were merged, and edge lengths were assigned as Euclidean distances between connected nodes. To improve network connectivity, disconnected fiber segments were repaired by connecting nearby endpoints separated by less than 10 μm and exhibiting an angular mismatch below 10°. Duplicate edges and self-loops were removed, and only the largest connected component of the reconstructed graph was retained for subsequent analyses. The final output consisted of a three-dimensional graph representation of the fiber network that can be modeled as described in **Discrete fiber model**.

### Network segmentation validation

To obtain an initial test of whether our segmentation procedure captured plausible network connectivity, we first analyzed the junction-degree distribution (node connectivity, defined as the number of elements that share a given node, denoted as z) of reconstructed collagen networks in our TFM experimental dataset of 35 images. Considering only network junctions (z>2), reconstructed networks were dominated by three-connected nodes (z=3; ~70%) and four-connected nodes (z=4;20%), with a smaller fraction of higher-order junctions (see histogram of network connectivity which includes all the node connectivities in the network **Extended Data Figure 1**b). This distribution is consistent with previous observations that collagen I networks have average connectivities between 3 and 4, and therefore remain below the Maxwell isostatic threshold in 3D (z=6)^45,46^. We then assessed whether fiber density, fiber location and orientation are also captured accurately with our segmentation workflow using ground-truth simulations.

We simulated synthetic ground truth (GT) networks that could be virtually corrupted by microscopy imaging noise, processed by our workflow and compared to (**Extended Data Figure 1**c). We first generated GT networks (**Extended Data Figure 1**c) using our previously described virtual fiber generator^40,41^, adjusted to produce average connectivities between 3 and 4 (**Extended Data Figure 1**d). We discretized fibers with minimum element length (MEL) of 0.36 µm, matching the experimental minimum pixel physical size. This choice was motivated by the fact that real image segmentations typically model fibers with element lengths larger than the pixel size to reduce computational cost. Therefore, we ensured that the resolution of the GT was at least as fine as the pixel size, allowing us to quantify the impact of coarser discretization during network recovery. Importantly, further refinement of the element size did not change the bulk mechanical behavior of the network, with clear deviations only happening for MELs > 2.52µm (**Extended Data Figure 1**e). We then generated synthetic confocal reflectance images by rasterizing the graph representation into a binary 3D skeleton image. Each graph element was sampled along its length according to the voxel size, and sampled points were assigned to the corresponding image voxels. The resulting binary skeleton was then convolved with an anisotropic 3D Gaussian point spread function to mimic the lower axial resolution of confocal microscopy (**Extended Data Figure 1**f,g). To match the experimental apparent fiber widths we first manually measured fiber lateral and axial intensity profiles in 20 randomly selected xy and xz image slices (**Extended Data Figure 1**f - green lines) and then adjusted the lateral and axial Gaussian widths accordingly (**Extended Data Figure 1**f - orange lines, **Extended Data Figure 1**g).

**Extended Data Figure 1:**
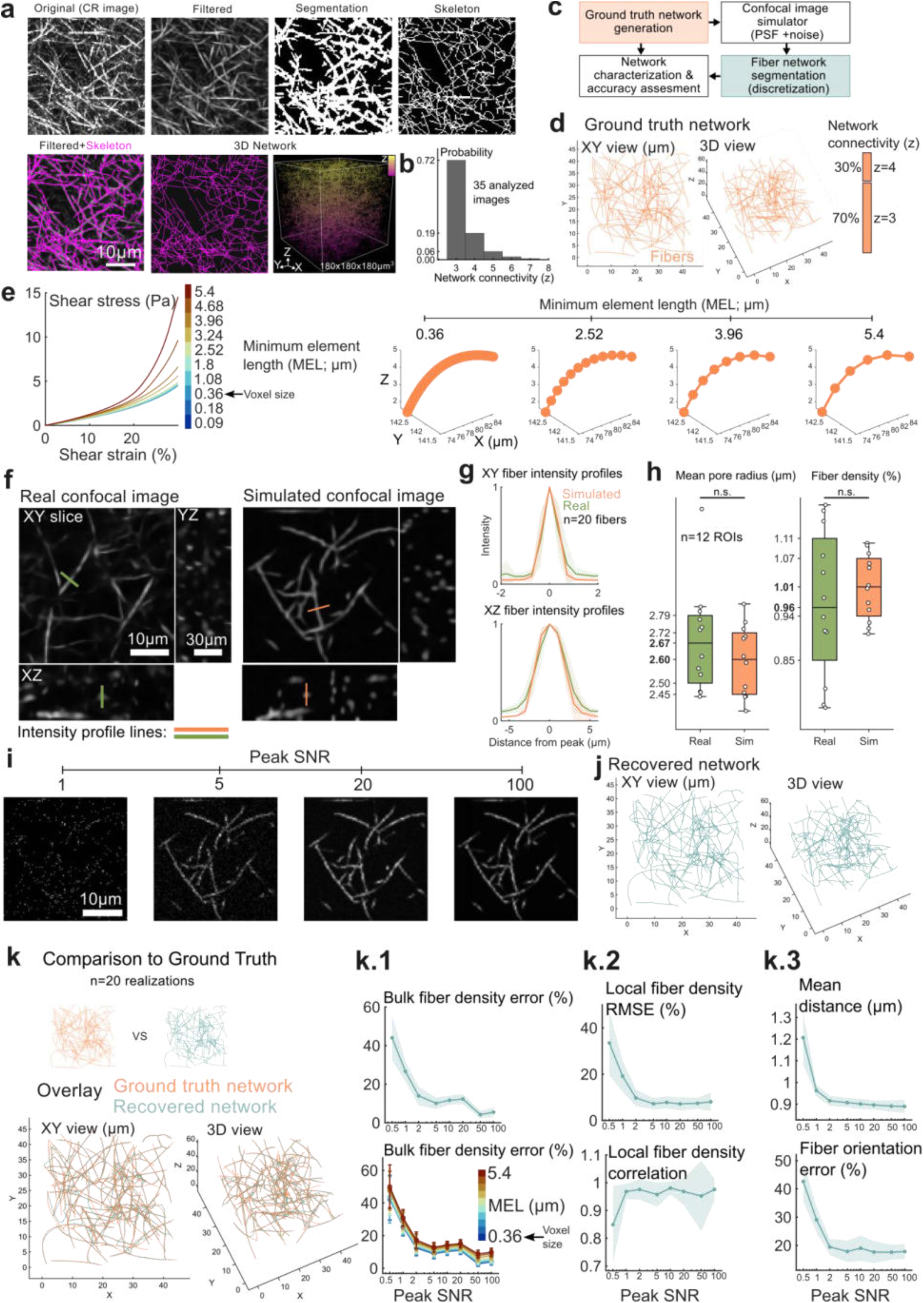
Validation of fiber segmentation algorithm. **a:** Overview of the image-processing workflow used to reconstruct collagen fiber networks from volumetric microscopy images. Top row: maximum intensity projection of a ROI of raw CR image, denoised (coherence-enhancing anisotropic diffusion filter) image, segmented binary fiber mask and resulting skeleton. Bottom row: overlay of the filtered (grayscale) and the skeleton (magenta) images, smoothed graph representation and a render of the full 3D (180×180×180µm3) extracted network colored by z-height. **b**: Network connectivity distribution of reconstructed collagen networks from the experimental TFM dataset. Only network junctions were considered (z>2). **c**: Schematic of the ground-truth validation workflow. Synthetic ground-truth networks were converted into simulated microscopy images, processed with the same segmentation workflow as experimental data and compared with the original ground-truth networks. **d**: Two views of a representative synthetic ground-truth network generated with an average connectivity between 3 and 4. **e**: Left: Sensitivity analysis of the influence of minimum element length (MEL) on the virtual shear response of the network. The experimental voxel size is indicated for reference. Right: 3D visual representation of the same fiber sampled with increasing MEL. Nodes are represented as dots and beam elements as straight lines. **f**: Representative experimental (left) and simulated (right) confocal reflectance images (one slice), including orthogonal views. Colored lines indicate representative intensity profiles measured across individual fibers. **g**: Mean lateral (top) and axial (bottom) fiber intensity profiles measured in simulated and experimental images. Profiles were centered at the position of maximum intensity. **h**: Comparison of simulated (green) and experimental (green) image morphology using an independent image-processing workflow. Mean pore radius (left) and fiber density (right) quantified in simulated (orange) and real image (green) regions. **i**: Visual comparison of the same simulated image slice shown in f), corrupted with increasing levels of peak shot-noise signal-to-noise ratio (peak SNR) noise. **j**: Representative recovered network obtained by applying the segmentation workflow to a simulated microscopy image originating from the GT network shown in d). **k**: Overlay of ground-truth and recovered networks, illustrating spatial agreement between both network representations (left) and error quantification of the recovered networks compared to ground truth as a function of microscopy image peak SNR (right panels): bulk fiber density for MEL = 2.52µm (k.1, top) and for a sweep of MELs (k.1, bottom); local fiber density RMSE (k.2, top); local fiber density correlation (k.2, bottom); mean distance between networks (k.3, top); fiber orientation error (k.3, bottom).

The resultant simulated images visually resembled the experimental ones (**Extended Data Figure 1**f) and matched the experimentally measured median pore radius and fiber density with errors below 5% and without statistically significant differences (**Extended Data Figure 1**h). These two quantities were measured using a simple image processing pipeline applied equally to real and simulated images. Although these metrics could have been computed using our own network reconstruction framework, doing so would have introduced a potential inverse crime, in which validation becomes biased toward the reconstruction method itself. Instead, we used an independent image-processing workflow to obtain an unbiased comparison. Briefly, images were first enhanced using a coherence filter, binarized using a fixed threshold, and objects smaller than 100 voxels were removed. The median value of the distance transform within the background phase was then computed as a proxy for the characteristic pore radius. Fiber density was estimated by skeletonizing the binary image and calculating the fraction of the volume of skeleton voxels relative to the total image volume.

Image noise was modeled as Poisson-distributed photon shot noise. The clean PSF-blurred image was scaled such that the brightest fiber voxel corresponded to a prescribed expected photon count. Noisy images were then generated by sampling each voxel intensity from a Poisson distribution. The corresponding peak shot-noise signal-to-noise ratio was defined as square root of the expected photon count (referred to as peak SNR in **Extended Data Figure 1**i,k.1-3). We used a voxel physical size matching that of the experimental images: 0.36×0.36×1µm.

We then applied our fiber segmentation algorithm to these synthetic images and compared the recovered network (**Extended Data Figure 1**j) to the ground truth. Qualitatively, our recovered networks closely overlapped with the GT networks (**Extended Data Figure 1**k). We then computed the following error metrics to assess the overall robustness of the network recovery against microscopy noise (20 realizations to account for stochasticity) using MEL=2.52µm (the largest value above which the mechanical response substantially deviates from that of the GT and sub-pixel MELs; **Extended Data Figure 1**e):

- **Bulk fiber density error** was computed as the relative difference between the recovered and ground-truth fiber densities. Fiber density was defined as the total fiber length per unit of volume: 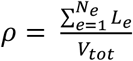, where *L*_*e*_ is the length of graph element *e, N*_*e*_ is the number of elements, and *V*_*tot*_ is the total analyzed domain volume. The error metric was computed as: 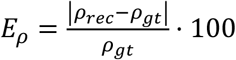, where *ρ*_*rec*_ and *ρ*_*gt*_denote the fiber densities of the ground-truth and recovered networks, respectively.
- **Local fiber density error** evaluated local variations in fiber density. The volume was partitioned into overlapping cubic sub-volumes. Within each cube *j*, graph elements were geometrically clipped to the cube boundaries, and the local fiber density, *ρ*_*j*_, was computed. The error metric was quantified using the normalized root mean squared error: 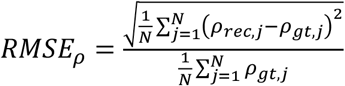, where *N* is the number of analyzed sub-cubes.
- **Local fiber density correlation** evaluated spatial variations in fiber density. The Pearson correlation coefficient between the ground-truth and recovered local density distributions was then calculated as: *r*_*ρ*_ = *corr*({*ρ*_*gt,j*_}, {*ρ*_*rec,j*_}).
- **Local fiber orientation error** was computed by partitioning the volume into overlapping cubic sub-volumes. Within each cube *j*, graph elements were geometrically clipped to the boundaries and a length-weighted orientation tensor, ***A***, was computed from the clipped segments. The relative error between the recovered and ground-truth local tensors was then calculated using the Frobenius norm and normalized by the Frobenius norm of the ground-truth tensor, 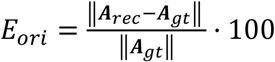. Cubes containing less than 1μm of ground-truth fiber length were excluded. The final local orientation error was summarized as the median value across all valid sub-volumes.
- **The mean distance** quantified how close the recovered nodes are to the ground truth nodes. For each node of the recovered network, *i*, the Euclidean distance to the closest node in the ground-truth network was computed as 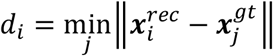, where 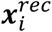 and 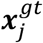 denote node coordinates in the recovered and ground-truth networks, respectively. The mean distance was then defined as the average of all *d*_*i*_.

For SNR>10 the bulk fiber density error remained below 10% while for lower SNR this metric increases to up to 40% (**Extended Data Figure 1**k.1, top). Varying MEL of the recovered networks also preserved this trend with slightly higher errors for lengths larger than the voxel size (**Extended Data Figure 1**k.1, bottom). This trend was also preserved when we locally (sub-cubes of 20×20×20 µm^3^ with 10µm spacing) analyzed the recovered networks with the local fiber density RMSE metric (**Extended Data Figure 1**k.2, top). Importantly, the local fiber density of the recovered networks remained highly correlated with that of the GT networks, with correlation coefficients close to 1 for peak SNR values above 0.5 (**Extended Data Figure 1**k.2, bottom). The mean distance between the recovered and GT networks was below 1µm for peak SNR values above 1 and plateaued at approximately 0.9 µm (**Extended Data Figure 1**k.3, top). Similarly, fiber orientation errors decreased with increasing SNR but did not fall below approximately 19%, even at the highest SNR values tested (**Extended Data Figure 1**k.3, bottom). This residual error is primarily attributed to the rasterization step used to convert the GT graph network into a voxelized microscopy image prior to image simulation. Although this step introduces discretization-related inaccuracies, it also reflects the finite spatial resolution inherent to microscopy imaging and therefore provides a realistic assessment of reconstruction performance. Consequently, these remaining errors can be interpreted as a limitation imposed by image discretization and resolution rather than by the segmentation algorithm itself.

While the present study aimed to reproduce imaging conditions representative of those typically encountered in 3DTFM, further improvements in reconstruction accuracy may be achievable through higher-resolution imaging approaches. Super-resolution techniques could reduce discretization artifacts and provide a more faithful representation of fiber architecture. Recently, a topology-preserving reconstruction framework for fibrous networks has been proposed^47^. Although this framework is highly attractive for small image volumes, we found that its computational cost increased rapidly for image stack sizes relevant to 3D TFM, requiring multiple weeks of computation on a high-performance computing cluster for a single image stack. Consequently, this approach could not be incorporated into the present study. Nevertheless, it represents a promising direction for future developments in DD-TFM.

### Virtual shear rheology and parameter calibration

Once the fiber network was extracted from confocal reflectance images, we performed virtual simple shear tests to ensure that the fiber-based model reproduced the macroscopic mechanical behavior of collagen gels. We followed the framework presented in Sanz-Herrera et al^40^. Macroscopic shear strain increments, which simulates the shear rheology test, were affinely transferred to the end nodes of the fiber network of the RVE. The resulting microscale fiber deformations and forces were computed using an updated Lagrangian finite element formulation in which elements are considered as Euler-Bernouilli beams. Macroscopic stresses were then obtained through Hill’s equality of energies, enforcing equality between the macroscopic strain energy density and the volume-averaged microscopic strain energy density. Details about the derivation of the stress tensor following this criterion can be found in^40^. This approach directly links the bulk mechanical response to the underlying fiber architecture and captures nonlinear mechanisms such as fiber straightening, alignment, buckling and strain stiffening^40,41^. We tested 4 sub-volumes of 75×75×75µm^3^ extracted from a segmented network image (orange line and shade in **Figure 1**d and **Figure 1**e).

*k*_*a*_ and *k*_*b*_ beam stiffnesses are assumed to be the same for all the elements of the model as a first approach. They were not derived directly from the apparent fiber diameter observed in the CR microscopy images as diffraction-limited optical microscopy does not have sufficient resolution. Instead, *k*_*a*_ and *k*_*b*_ were independently fitted to match the macroscopic mechanical response of collagen gels obtained from the shear rheology measurements. This modeling choice is motivated by the multiscale and heterogeneous nature of collagen fibers, which often appear as dense bundles of nanofibrils with irregular packing^45,48^. As a result, treating the fibers as solid, homogeneous cylinders, where cross-sectional area, *A*_*f*_, and second moment of inertia, *I*_*f*_, are directly computed from the observed diameter (Equation 1, Equation 2), can significantly overestimate their load-bearing capacity. This issue is analogous to modeling a structural cable: although it may appear cylindrical, it has an internal structure (number of filaments, braiding pattern, air gaps) that makes it not mechanically behave like a solid rod of the same outer diameter. In real structural cables, the Young’s modulus can be associated with the material composing the filaments (e.g., steel), but the cross-sectional area and moments of inertia are treated as *effective* properties that account for the complex internal organization. Therefore, in our model, we treat both *k*_*a*_ and *k*_*b*_ as *effective* and independently fitted parameters, without directly linking them to a physically measurable diameter or modulus. Nevertheless, the values obtained for this paper (*k*_*a*_=17 MPa·µm^2^; *k*_*b*_=7 kPa· µm^4^) are realistic. Theoretically, they would yield a fiber Young’s modulus of 3.29GPa and a fiber diameter of 81.2nm, both within the order of magnitude of what was reported via AFM measurements (Young’s moduli ranging ~0.25-11GPa and diameters from 50-200nm)^32,33^. In any case, both *k*_*a*_ and *k*_*b*_ are physically measurable quantities that could be obtained directly through single fiber tests. This approach allows us to reflect the multiscale and anisotropic behavior of the fibrous collagen matrix while maintaining consistency with macroscopic experimental data. *k*_*t*_ can be expressed as a function of *k*_*b*_ and *v* (Equation 3). In this study we assumed *v*=0.3. Nevertheless, prior works have shown that torsional contributions have little impact on the total accumulated matrix energy^40,41,49^.

### Fiber displacement measurement

Matrix displacements were estimated using the image-processing and registration workflow implemented in TFMLAB^50^. Briefly, relaxed and stressed image stacks were first filtered and corrected for rigid shifts. Non-rigid displacement fields were then obtained using a free-form deformation (FFD) image-registration approach^15,51^. In this method, the relaxed and stressed image stacks are treated as moving and fixed images, respectively, and the displacement field is obtained by deforming the relaxed image until it optimally matches the stressed image. The deformation is represented by a smooth B-spline transformation defined on a regular grid of control points. During registration, the positions of these control points are iteratively updated to minimize an image similarity metric, while regularization ensures a spatially smooth deformation field. The resulting continuous (voxel-based) three-dimensional displacement field was subsequently interpolated at the nodes of the reconstructed fiber network.

### Force recovery methods for DD-TFM

Let *N* be the number of nodes of the network. Each node *i* has 6 degrees of freedom (DOFs): 3 translational and 3 rotational. ***u***_*i*_∈ℝ^3^, ***θ***_*i*_∈ℝ^3^, ***f***_*i*_∈ℝ^3^ and ***m***_*i*_∈ℝ^3^ are the translations, rotations, nodal forces and nodal moments of node *i*, respectively. The global displacement and force vectors, ***U*** and ***F***, are defined as 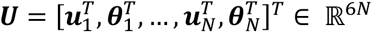 and as 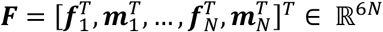, respectively. In order to capture geometrical nonlinearities, we follow an updated Lagrangian FEM approach^40^:

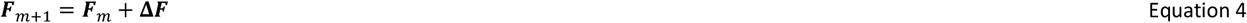

Such that our beam network discretized elasticity problem turns into the following algebraic system:

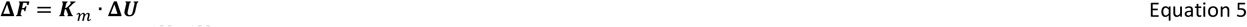

Where ***K***_*m*_∈ℝ^6*Nx*6*N*^ is the updated tangent stiffness matrix, constructed in the current configuration *m* after updating nodal coordinates, from element contributions and incorporating both translational and rotational stiffness terms. Forces can be recovered with the forward method or with inverse method. In this paper we test two inverse methods: a classic Tikhonov regularization approach and a hybrid version of our physics-based inverse method (*HPBIM*^34^) in which, in addition to Tikhonov regularization, equilibrium of forces is imposed in the hydrogel domain^35^. These methods are referred to as *Classic* and *HPBIM*, respectively.

*Forward method*. The global displacement vector for the forward method, ***U***_*fwd*_, is only partially known since it includes translations and rotations. Let *ℸ* denote the set of translational DOFs and ℛ the rotational DOFs. We can write the displacement vector as:

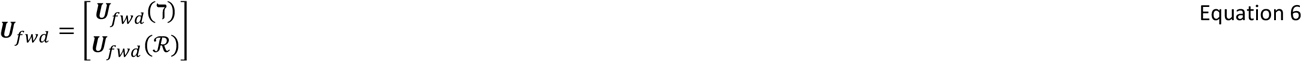

***U***_*fwd*_(*ℸ*) is known, as it is equal to the measured displacement field interpolated to the fiber network nodes, ***u***_*meas*_. To account for geometric nonlinearities, we divide the displacement vector into *M* sufficiently small incremental configurations **Δ*U***_*fwd*_(*ℸ*) = ***U***_*fwd*_(*ℸ*)/*M*, such that the small displacement assumption holds for *m* → *m* + 1 configurations. We then rearrange the linear system (Equation 5) accordingly:

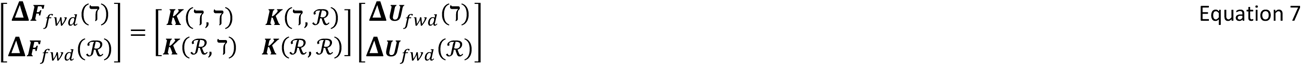

Note that subindex *m* has been omitted in stiffness matrices to simplify the notation. Assuming that no external moments are applied (i.e., ***F***_*fwd*_(ℛ) = 0), we solve the reduced system:

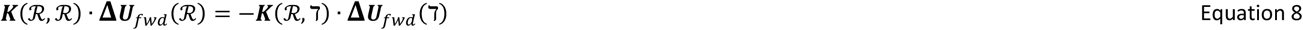

This equation is solved through Matlab’s function `mldivide`, yielding the unknown nodal rotations **Δ*U***_*fwd*_(ℛ), which are concatenated with **Δ*U***_*fwd*_(*ℸ*) to form the full global displacement vector **Δ*U***_*fwd*_. Finally, the corresponding forces, ***F***_*fwd*_, is computed iteratively via Equation 4 and Equation 5. Because **Δ*U***_*fwd*_ contains measurement noise, we make use of the so-called inverse methods.

*Inverse methods*. Distinguishing two problem domains that include all 6 DOFs, i.e., the DOFs corresponding to the fiber network nodes closest to the surface of the cell and at the boundaries of the ROI, A, and the DOFs of the hydrogel internal nodes, B (see **Extended Data Figure 2**a,b), we can reorder Equation 5 as:

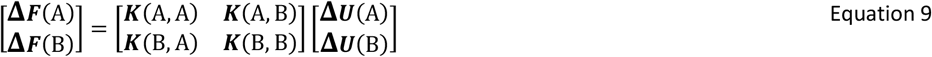

Inverse methods (*Classic* and *HPBIM*) search for an alternative displacement vector, **Δ*U***_*inv*_, that is as close as possible to **Δ*U***_*fwd*_ with a penalty for high force vector norms through Tikhonov regularization^35^:

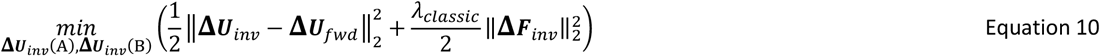

*HPBIM* additionally imposes the fulfillment of the equilibrium of forces in the hydrogel domain, B^34,35,52^. These ideas can be mathematically written as:

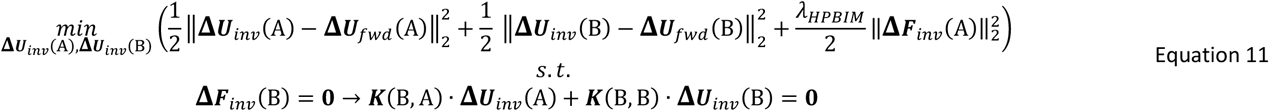

The equilibrium constraint can be included as a Lagrange’s multiplier ***η***, resulting in:

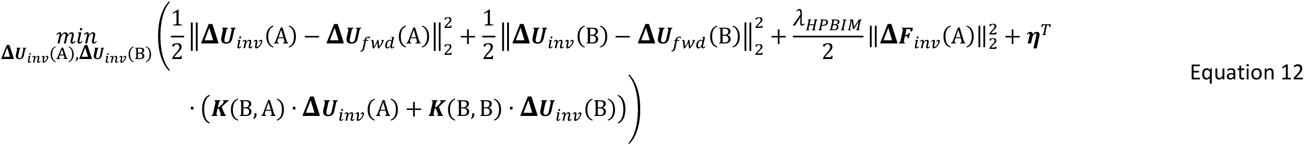

Note that, unlike in the *Classic* formulation (Equation 10), equilibrium is fulfilled even for non-zero values of *λ*_*HPBIM*_. The minimum of Equation 12 can be analytically derived as:

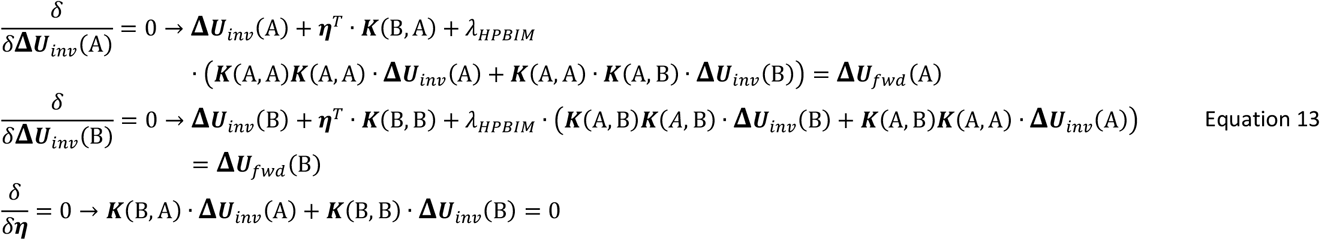

This system of linear equations can be written in matrix form as:

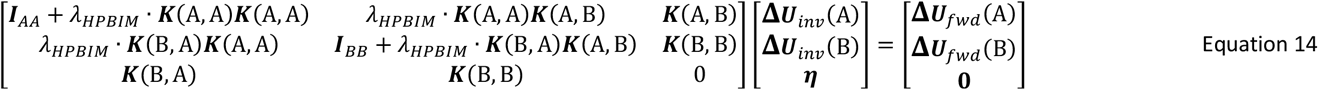

This ***a*** · ***x*** = ***b*** linear system is solved through Matlab’s function `mldivide` to obtain **Δ*U***_*inv*_. Note that the *Classic* method does not use the third term of Equation 12 and therefore the matrices of Equation 14 loose their third row. Moreover, it regularizes forces both in A and B domains. Finally, ***F***_*inv*_ is then computed iteratively using Equation 4 and Equation 5. We performed a sensitivity analysis to determine the number of incremental configurations (*M*) used to recover forces (**Extended Data Figure 2**c-top). The norm of the recovered force field stabilized after dividing the displacement field into ~35 configurations, with variations of only a few pN for larger values of *M*. On a 64-bit Windows 11 Enterprise workstation equipped with an 11th Gen Intel Core i7-11800H CPU running at 2.30 GHz and 32 GB of RAM, each configuration required ~4 minutes (see **Extended Data Figure 2**c-bottom). To ensure stable force recovery with a reasonable computational cost, we used *M* = 40 configurations throughout this study (see **Supplementary Videos 1, 2**). The value of *λ*_*Classic*_ or *λ*_*HPBIM*_ was obtained using the L-curve method. Briefly, force reconstruction was performed over a range of *λ* values from 10^−8^ to 10^−1^ and ‖***F***_*inv*_(A)‖_2_ was plotted against ‖***U***_*inv*_ − ***U***_*fwd*_‖_2_. The optimal value was defined as the point of maximum curvature of the resulting L-curve, which simultaneously minimizes ‖***F***_*inv*_(A)‖_2_ and ‖***U***_*inv*_ − ***U***_*fwd*_‖_2_. For the in silico simulations, this procedure was performed independently for each noise level and force recovery method. For the experimental dataset (35 cells), we only used *HPBIM*. All samples were acquired under the same imaging and experimental conditions and were therefore expected to have comparable displacement noise levels. We determined the optimal *λ*_*HPBIM*_ value for one representative cell and used this value (*λ*_*HPBIM*_=7·10^−5^) for the remaining experimental samples **Extended Data Figure 2**d,e. Moreover, we verified that the recovered forces in the unconstrained *B* domain were negligible compared with those in the *A* domain. Across the 35 experimental cells, the median force norm in the *B* domain was on the order of 10^−8^ pN, whereas the median force norm in the *A* domain was on the order of 10^3^ pN (**Extended Data Figure 2**f). Global force balance was also satisfied with high precision, with the summed *x*-, *y*- and *z*-components of the forces in the *A* domain remaining on the order of 10^−7^ pN (**Extended Data Figure 2**g). Moreover, we verified the numerical precision of the solution to the ***a*** · ***x*** = ***b*** linear system of Equation 14 obtained with MATLAB’s `mldivide` by computing the relative residual, ‖***ax*** − ***b***‖_2_/‖***b***‖_2_, at each of the 40 incremental configurations for every cell. The median, 95th percentile and maximum residuals remained ~10^−6^, verifying stable numerical solution of the inverse problem (**Extended Data Figure 2**h).

**Extended Data Figure 2:**
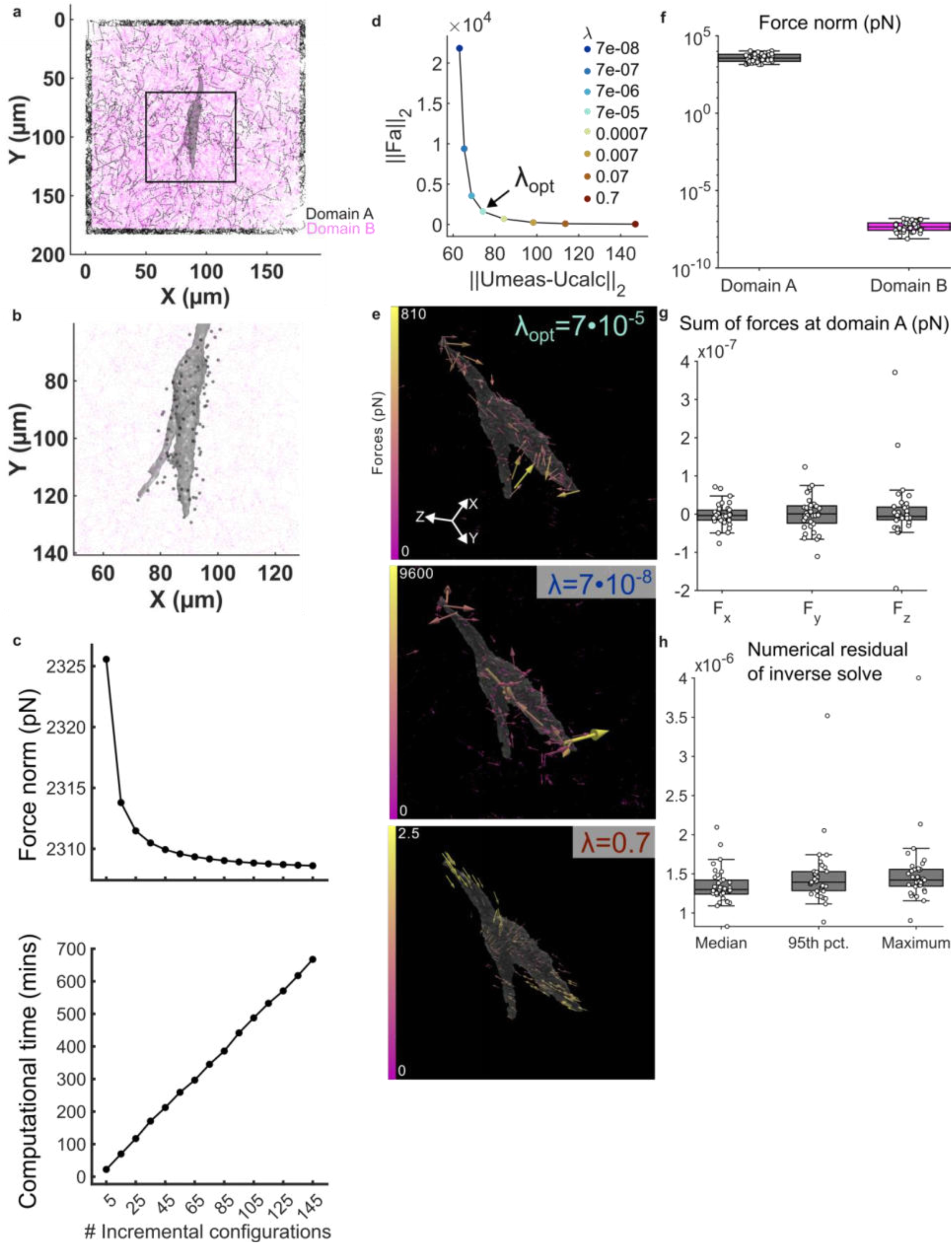
**a**: 3D view of the two computational domains used for HPBIM. Domain A are the network nodes closest to the surface of the cell and the boundaries of the ROI, and B are the hydrogel internal nodes. **b:** Close up into the black frame marked in a) with the nodes closest to the surface of the cell enhanced for visibility. **c**: Sensitivity of recovered norm of the force vector (top) and computational cost (bottom) to the number of incremental deformation configurations used in HPBIM. **d:** Representative L-curve used to select the regularization parameter for HPBIM. The optimal value was defined as the point of maximum curvature (arrow; 7e-5). **e**: Visual comparison of the resultant force fields with different values of λ, including the optimal value (top), and underestimated one (middle), and an overestimated one (bottom). **f:** Comparison of recovered force norms in both domains, showing that forces recovered in the B domain were negligible. **g:** Global force-balance check for experimental DD-TFM reconstructions. Summed force components in the A domain remained close to zero in the x-,y- and z-directions. **h:** Numerical residuals of the HPBIM linear solve, confirming stable solution of the inverse problem with MATLAB’s mldivide for all 35 cells of the dataset. Each point represents the median (left), 95^th^ percentile (middle) or maximum (right) residual (across the 40 incremental configurations) of a cell.

We compared CT and Blebb experimental conditions by means of metrics computed at the network nodes closest to the cell surface, *C*: strain energy, defined as: 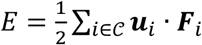, total force, defined as: *F*_*tot*_ = ∑_*i*∈*C*_ ‖***F*** _*i*_ ‖, and total contractility, defined as 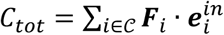, with 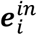 being the inward unit vector from the node position ***x***_*i*_ to the cell centroid 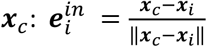. For each metric the CT and Blebb distributions were compared using MATLAB’s `ranksum` function, corresponding to a two-sided Wilcoxon rank-sum test. Group summaries were displayed as mean ± SEM, with individual cells shown as overlaid jittered points (**Figure 2**b). No paired structure was assumed between conditions. Additionally, we computed the mean, median, standard deviation and 98th-percentile force from the distribution of force magnitudes, {‖***F***_*i*_‖}_*i*∈*C*_ (**Extended Data Figure 4**a-d).

### Fiber stresses and energies in DD-TFM

After recovery of nodal displacements with *HPBIM*, fiber stresses and strain-energy contributions were calculated at the element level for each incremental deformation step. Each collagen fiber element was treated as a two-node Euler-Bernoulli beam with six degrees of freedom per node. For each element, a local coordinate system was defined from the current element orientation: the local axial direction was given by the vector connecting the two element nodes, and two orthogonal transverse directions were used to transform the 12-component nodal displacement vector from the global coordinate system to the local element coordinate system. The local beam stiffness matrix was assembled using the calibrated axial stiffness, *k*_*a*_, bending stiffness, *k*_*b*_, and torsional stiffness, *k*_*t*_. For each deformation increment, the local generalized force increment of element *e* was computed as:

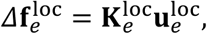

where 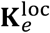 is the local beam stiffness matrix and 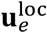 is the local element displacement vector. Generalized element forces were accumulated over deformation increments to obtain the current local force vector, containing axial forces, shear forces, torsional moments and bending moments at both element nodes. Axial, bending and torsional energy contributions were then calculated from the corresponding work-conjugate local force-displacement products,

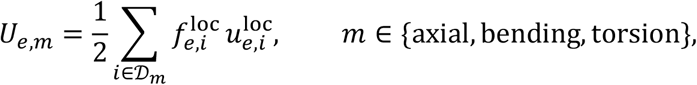

where *D*_*m*_ denotes the local degrees of freedom associated with each deformation mode. The axial contribution was computed from the axial force and axial displacement components, the torsional contribution from torsional moment and rotation components, and the bending contribution from the transverse force, transverse displacement, bending moment and rotational components. Elemental energy contributions were accumulated over all deformation increments and normalized by the analyzed network volume.

Fiber stresses were calculated from the accumulated local generalized forces. Axial stress was computed as 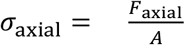, where *F*_axial_ is the local axial force and *A* is the effective fiber cross-sectional area, 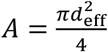. The effective diameter, *d*_eff_, was obtained from the calibrated axial and bending stiffnesses as 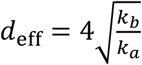. Bending stress was estimated from the maximum absolute bending moment at the element ends as 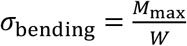, where *M*_max_ is the largest absolute bending moment around the two local transverse axes and *W* is the section modulus of a circular cross-section, 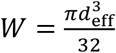. The total fiber stress was defined as the maximum normal stress obtained by superimposing axial and bending contributions,

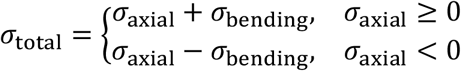

This sign convention preserves tensile and compressive axial loading while accounting for the maximum additional normal stress induced by bending.

To analyze how fiber stresses propagated away from the cell, we computed a total-stress profile as a function of distance to the cell surface. For each fiber element, the element centroid was calculated from the final *HPBIM*-recovered configuration, and its distance to the cell surface was approximated as the minimum Euclidean distance to the vertices of the segmented cell surface. Elements were assigned to 1 µm-wide distance bins from 0 to 40 µm. Within each bin, total stress (*σ*_total_) was averaged using a fiber-length-weighted mean, so that longer elements contributed proportionally more to the bin value. The resulting stress profile for each cell was smoothed with a 5 µm moving window and then averaged across cells within each condition for visualization (see **Figure 2**c).

To analyze the evolution of axial and bending energy during matrix deformation, we local regional energy metrics at each incremental *HPBIM* deformation step. For each fiber element, the centroid was calculated and its distance to the cell surface was approximated as the minimum Euclidean distance to the vertices of the segmented cell surface. Elements located within 5 µm of the cell surface were selected, and the median axial and bending energy across these elements was computed at each deformation step. This yielded one axial-energy and one bending-energy trajectory per cell. For visualization, trajectories were averaged across cells within each condition, with the incremental deformation step used as a pseudotime coordinate (see **Figure 2**e). Final axial and bending energies were compared between conditions using the last deformation step (see **Figure 2**f).

### In silico ground truth simulations and validation

To evaluate which method is more robust against measurement noise in this data-driven framework, we simulated a contracting cell embedded in a real collagen matrix (**Extended Data Figure 3**a,b) generating ground truth (GT) matrix displacements and cell forces (**Extended Data Figure 3**e-g). After acquiring the microscopy image of a cell embedded in collagen a cell binary mask was obtained by Otsu’s thresholding using Matlab’s function `imbinarize`.

**Extended Data Figure 3:**
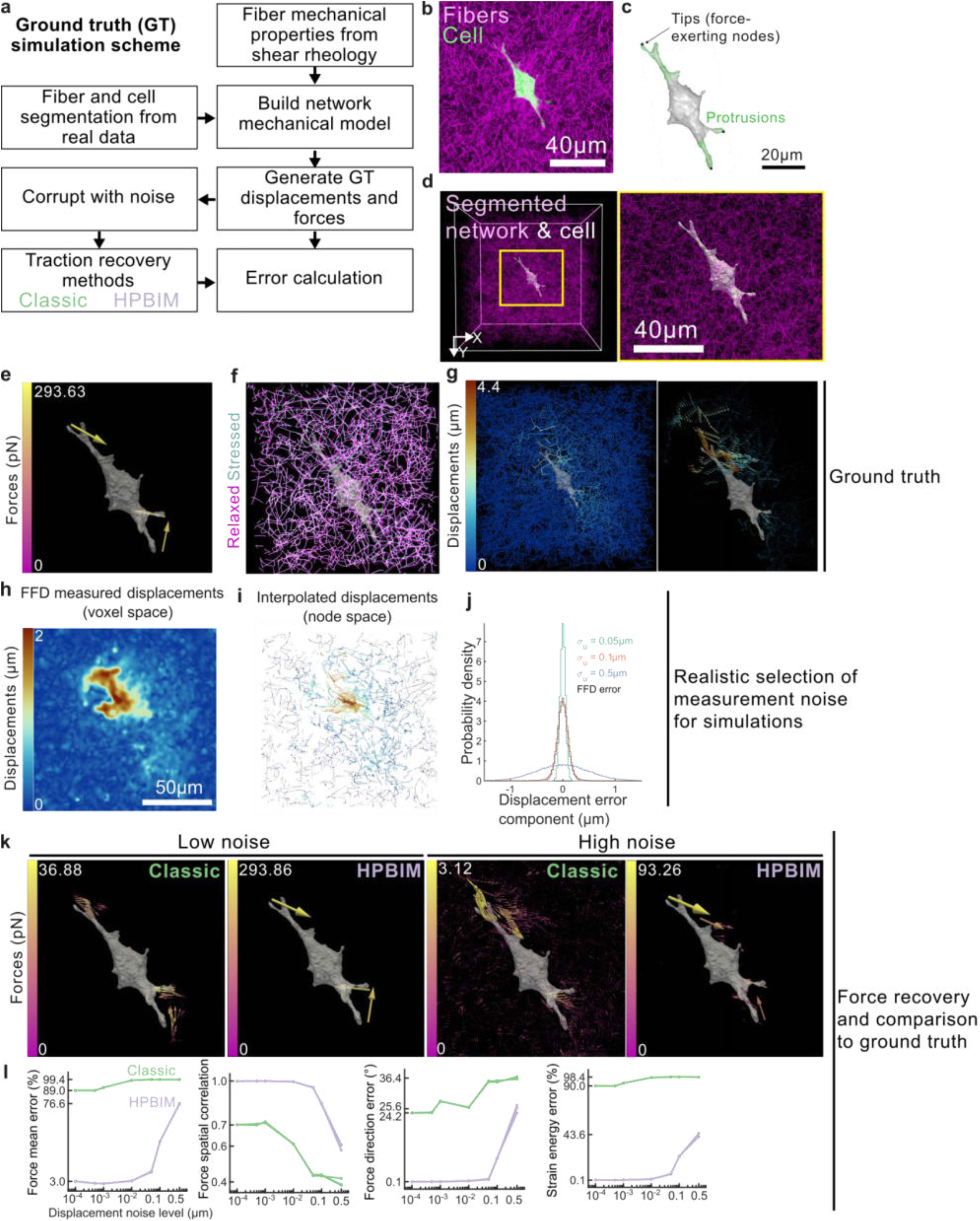
In silico validation of DD-TFM force recovery. **a:** Schematic of the in silico validation workflow. Fiber network and cell geometries were segmented from real microscopy images, and the mechanical parameters extracted from shear rheology were applied to the network mechanical model in order to represent a simulated contracting cell in a collagen fiber network. Ground-truth (GT) displacements and forces are generated; controlled Gaussian noise was then added to the displacement field, and forces were recovered using the Classic and HPBIM inverse methods and compared to the GT using error metrics. **b:** Maximum intenstity projection of a real image of a cell (CellTracker; green) embedded in collagen (CR, magenta). **c**: Segmented 3D cell morphology with detected protrusion and tip nodes in green and black, respectively. **d**: 3D view of the simulated model showing the cell embedded in a collagen network (left), and a closeup of the region marked in yellow (right). **e:** 3D render of the GT force field. Force vector arrows originate from fiber nodes. f: overlay of the relaxed (magenta) and stressed network (after applying GT displacements; blue). g: GT displacements represented as colored-by-magnitude fibers (cross-section; left) and as an arrow vector field (right) around the cell. **h**: Slice of the measured displacement field magnitude by means of FFD applied to simulated CR images. Colorbar is saturated to ease visualization. **i**: Interpolated measured displacement vectors from the image grid space to the fiber network nodal space. Colorbar is shared with h. **j:** Zero-mean Gaussian noise distributions with standard deviations from 0.05 to 0.5 µm (colors). The experimental FFD error distribution (black) was approximately Gaussian with standard deviation σ_u_ ≈0.1µm. **k**: Representative examples of the force recovery under low noise (noise level = 10^−4^) and high noise (noise level = 0.5) for Classic (green) and HPBIM (purple) methods. **l:** Validation metrics comparing Classic (green lines) and HPBIM (purple lines) force recovery as a function of displacement noise level, including force mean error, force spatial correlation, force direction error and strain energy error. The errors of each method are depicted by 3 lines, corresponding to 3 independent realizations of noise.

Then, a sphere of maximum radius was fitted within the cell volume. Every mask point located at a distance (with respect to the sphere center) equal to or higher than 1.5 times the sphere’s radius was considered a cell protrusion. The principal direction of each protrusion was obtained by calculating the eigenvectors of the protrusion mask with Matlab’s function `regionprops3`. The protrusion points with largest distance to the surface of the sphere was taken as the protrusion tips (**Extended Data Figure 3**c). We then positioned the cell segmentation at the center of the segmented fiber network (**Extended Data Figure 3**d) and an inward displacement of ~4 microns in the direction of the protrusions’ principal direction was defined at the closest fiber point to each protrusion tip. By imposing these displacements in the fiber model, ground truth displacements and forces were obtained by applying Equation 7 and Equation 8 (**Extended Data Figure 3**e-g).

To simulate realistic force recovery scenarios where the measured displacement contains noise, we added synthetic Gaussian noise to the ground truth displacement field. For each noise level (σ_u_), the corrupted displacement field was generated as:

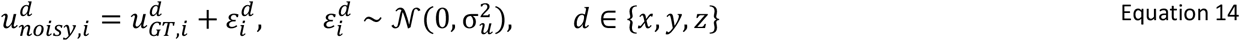

Where 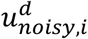 is the noise-corrupted displacement used as input for force recovery, 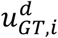 is the ground-truth displacement component *d* at network node *i*, and 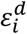 is an independent random displacement error drawn from a zero-mean Gaussian distribution. The parameter σ_u_ defines the standard deviation of the added noise and therefore corresponds to the prescribed noise level.

To select realistic noise levels close to those expected from real measurements, we first evaluated the displacement measurement errors obtained from FFD-based non-rigid registration (see **Fiber displacement measurement**). We converted our relaxed and stressed networks (with fibers displaced with the known ground truth displacements, see **Extended Data Figure 3**f) into synthetic image pairs (generated as described in **Network segmentation validation**). We then registered those images with FFD to obtain measured displacements on the image grid (**Extended Data Figure 3**h), which were then interpolated back to the relaxed fiber-network nodes (**Extended Data Figure 3**i). Displacement measurement errors were computed at each selected node as the difference between the FFD-measured and ground-truth displacement vectors. To focus on mechanically relevant regions, this error was evaluated for nodes whose ground-truth displacement magnitude was above the 75th percentile. The x, y and z components of the displacement error vectors were then pooled, yielding an approximately Gaussian component-wise error distribution with standard deviation σ_u_ ≈0.1µm (see **Extended Data Figure 3**j). Because improved fiber image acquisition and segmentation may further reduce displacement noise (see discussion in **Network segmentation validation**), and to assess the robustness of force recovery more generally, we tested a range of imposed displacement-noise amplitudes spanning values below and above the measured level: σ_u_ = 10^−4^ to 5·10^−1^ µm. We then systematically corrupted the GT displacement fields with these different levels of noise (3 realizations per noise level) and recovered forces with *Classic* and *HPBIM* (see **Recovery of cell forces**).

To evaluate the accuracy of the reconstructed force fields using *Classic* or *PBNIM*, the metrics shown in **Table 1** were computed at the network nodes closest to the cell surface, *C*, and plotted as a function of displacement noise in **Extended Data Figure 3**l.

**Table 1:**
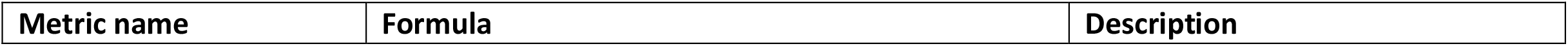
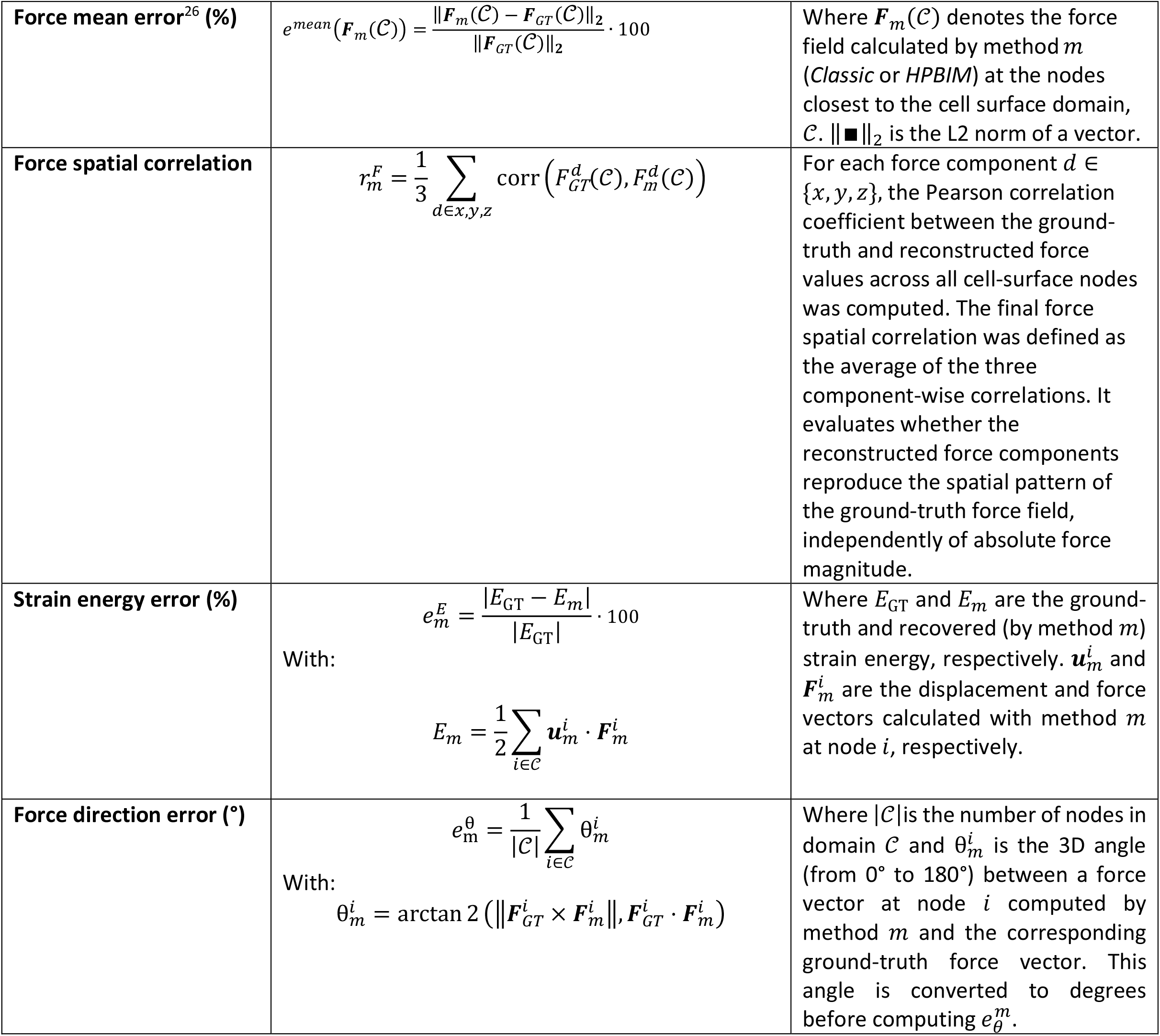
Metrics used to evaluate the accuracy of force recovery methods in DDTFM.

Qualitatively, *Classic* recovered smooth force patterns that strongly underestimated the GT force magnitude and introduced unrealistic non-zero forces away from the cell (**Extended Data Figure 3**k). In contrast, *HPBIM* more accurately preserved force magnitudes and consistently recovered clear pulling patterns, even when displacement fields were heavily corrupted (**Extended Data Figure 3**k). Quantitatively, all error metrics report an overall superiority of *HPBIM* over *Classic*. In the presence of noise, *Classic* showed mean force errors of ~90%, whereas *HPBIM* remained below 40% except at the highest noise level, where the error reached ~76% (**Extended Data Figure 3**l.1). Despite these force magnitude errors, *HPBIM* recovered spatial distribution of forces with high fidelity, with force spatial correlation values between 0.8 and 1, whereas *Classic* remained below 0.7 for most noise cases (**Extended Data Figure 3**l.2). *HPBIM* captured force vector directions also more accurately, with errors below 10°, compared to errors >25° for the *Classic* in most noise cases (**Extended Data Figure 3**l.3). Strain energy was also more accurately captured by *HPBIM*, with errors around 0.1% for most noise cases and a maximum error of 43% in the noisiest case, while *Classic* shows errors above 90% (**Extended Data Figure 3**l.4). These findings highlight a key limitation of 3D TFM: noise in displacement measurements propagate strongly into force calculation, consistent with what has been observed in conventional, continuum-based 3D TFM^12,22,24,26,34,52^. Given the inevitably noisy measurements due to image acquisition by CR microscopy and network segmentation, these simulations conclude that *HPBIM* provides the most reliable force recovery in real DD-TFM.

### Continuum 3D TFM

#### Continuum version of HPBIM for 3D traction reconstruction

Earlier, we proposed a continuum traction recovery method for 3D TFM that, as *HPBIM* does, searches for a new displacement field, which must optimally resemble the noisy measured one while simultaneously fulfilling the equilibrium condition within the ECM’s analogue medium (collagen hydrogel)^35^. However, this method was formulated following a Finite Element variational approach by means of the Principle of Virtual Work. In this sense, the problem is treated as a Lagrange optimization procedure, where the difference between the new and measured displacement fields is minimized and the equilibrium condition is inserted into the objective function by means of a Lagrange multiplier field. Moreover, the objective function also includes a Tikhonov regularization term, controlled by a scalar parameter, which represents the degree of penalization applied to the values of surface and volume forces during the minimization process. This forces the solution to deviate to a greater extent from the input displacements relative to those obtained without regularization, ensuring a certain degree of smoothness. In this method, only forces acting on the cell surface are regularized, since forces within the hydrogel’s domain are kept nil by the enforcement of the equilibrium condition. Similar to Equation 11, the mathematical formulation of the problem described above can be stated as follows:

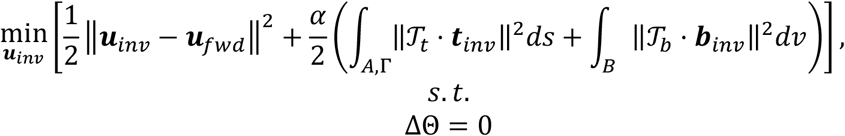

in which ***t***_*inv*_ and ***b***_*inv*_ represent, respectively, the surface and volume (or body) forces acting on surfaces A and the inner ECM’s domain B (the definition of these domains is analogous to the one described for the fiber nodes in **Extended Data Figure 2**a). *T*_*t*_(*x, y, z*) and *T*_*b*_(*x, y, z*) are scalar fields bounded in the interval [0,1] which control the contributions to the objective function of the corresponding forces within the referenced regions. ΔΘ represents the equilibrium constraint enforced within the ECM’s domain. This problem features both material (hyperelasticity) and geometrical (large deformations) nonlinearities, hence its solution requires its linearization in the context of a Newton-Raphson iterative procedure. In addition, the featured variables are in turn discretized following a Finite Element approach, which transforms the regularization term into the following manner:

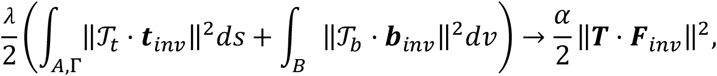

being ***F***_*inv*_ the array containing both surface and body forces acting on each degree of freedom associated with the nodes that compose the finite element mesh. Furthermore, ***T*** represents the discretization of the scalar fields *T*_*t*_(*x, y, z*) and *T*_*b*_(*x, y, z*), constituting a diagonal matrix that contains the weights corresponding to the contribution to the objective function of the forces acting on each degree of freedom of the mesh. It features only values of 1 and 0 distributed throughout its diagonal components, the former corresponding to the forces set to be regularized. In this case, only the degrees of freedom associated with nodes contained in the surface region *A* are regularized (receive a unit value). The now discretized constraint **Θ** = **0** is therefore applied via the discretized Lagrange multiplier field ***η***, resulting in the term **Θ** · **η** being introduced into the objective function (all variables are assumed to have been discretized at this point),

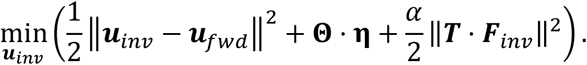

The Lagrange multiplier field ***η*** so happens to represent the displacements caused by the virtual forces resulting from the discrepancy between noisy (forward) and calculated (inverse) displacements, mathematically occupying the place of the virtual displacements in the equilibrium constraint. To find the system of equations required to retrieve ***u***_*inv*_ and ***η***, the objective function is differentiated with respect to both variables,

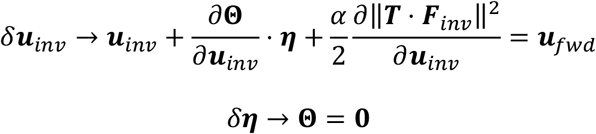

Within the Newton-Raphson linearization, *∂***Θ**/*∂****u***_*inv*_ corresponds to the tangent stiffness matrix ***K*** at a certain step within the iterative process. After the development of the other terms^35^, the final discretized system of equations receives the expression:

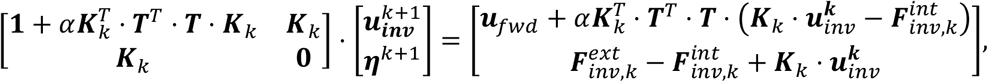

where **1** is the identity tensor and 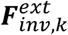 and 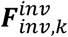 are the arrays containing the nodal forces (for each degree of freedom) respectively corresponding to external and internal forces. Known variables are evaluated at the current iteration step *k* (with the first iteration the initial guess), and the unknown variables are computed at the next step *k* + 1. This process is executed until the relative difference between the computed variables evaluated at *k* + 1 and at *k* is sufficiently small with respect to the established convergence criterion.

#### Nonlinear hyperelastic model

The continuous model employed to characterize the mechanical behavior of the collagen hydrogel utilized in the *in vitro* 3D TFM experiments is nonlinear hyperelastic, based on the one presented by Steinwachs et al.^24^. The latter gives a definition of the strain energy density function which contemplates the behavior of a particular collagen fiber. To find the macroscopic bulk’s behavior, the contribution of one fiber needs to be homogenized by considering all possible directions, which we did by means of performing the Gauss-Legendre quadrature over the unit sphere. A fiber is considered to be able to deform within three distinct regimes: (i) the fiber buckles when subjected to compression forces, its stiffness exponentially decaying to zero the more its ends are brought near each other; (ii) the fiber, being in a “collapsed” state, is straightened by means of axial forces, its stiffness remaining constant during the whole process until the distance between its ends matches its natural length; and (iiii) the fiber is extended beyond its natural length, its stiffness growing exponentially the further its ends are separated from each other. However, we found that a modified version of this model better fitted their experimental constitutive data, i.e., the shear rheology curves of the collagen hydrogels. Not only that, but this modification also more accurately replicates the synthetic shear rheology curves that we obtain with the multiscale fibered matrix generator model they developed and described in Sanz-Herrera et al.^40^. The modified model differs from the original in that the behavior corresponding to states of deformation (ii) and (iii) are interchanged in the former with respect to the latter. In this way, the strain energy density function is defined for a single fiber via its second derivative with respect to the macroscopic deformation measure, i.e., via its mechanical stiffness,

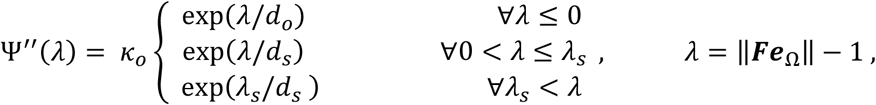

in which *κ*_*o*_ is the stiffness-analogue parameter, *d*_*o*_ is a dimensionless parameter controlling stiffness decay during buckling of the fiber, *d*_*s*_ is a dimensionless parameter associated with strain-stiffening behavior and *λ*_*s*_ is the dimensionless parameter that defines the extension of the interval of exponential strain-stiffening behavior. ***F*** represents the deformation gradient, ***e***_Ω_ is the unit vector defining the orientation of the fiber and *λ* is its macroscopic deformation, defined as the relative variation in length of a fiber with respect to its natural length. Then, the mechanical stress experienced by the fiber in terms of the macroscopic deformation is defined as,

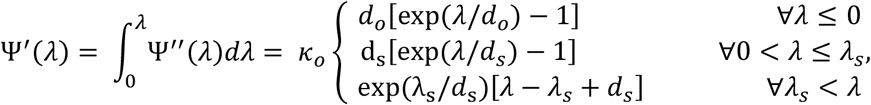

where integration starts at zero because the material is assumed to not be pre-stressed. The First Piola-Kirchhoff tensor, which embodies the constitutive relation in mixed configuration, can be obtained as follows:

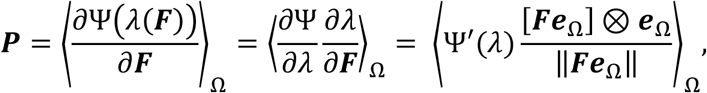

in which the angle brackets indicate averaging of the contributions of fibers over all possible directions Ω within the unit sphere. Then the Cauchy stress tensor, which corresponds to the constitutive relation in spatial configuration, can be easily obtained by means of a push-forward operation applied to 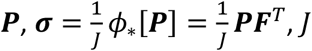 corresponding to the determinant of the deformation gradient tensor. The averaging procedure has been performed by means of the Gauss-Legendre quadrature, for which further details, as well as information regarding the computational implementation of the model within the Finite Element framework, can be found in Apolinar-Fernández et al.^27^.

To verify that the hyperelastic model can reproduce the modeled material’s behavior, the authors fitted the governing parameters of the model to experimental shear rheology data obtained from applying simple shear experiments on collagen hydrogels in the laboratory. In particular, the hydrogel was subjected to the action of a rotational rheometer of cone and plate, and as a result the stress-strain relationship of the material was measured. Concerning a simple shear state of deformation, the deformation gradient receives the expression,

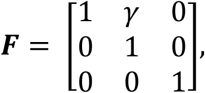

where *γ* denotes the engineering shear strain. The shear stress measured during the experiment corresponds to the shear component of the First Piola-Kirchhoff stress tensor (nominal stress), since the shear stress is computed from the retrieved torque together with the geometrical characteristics of the test in the nominal (undeformed) configuration. By means of the definition of the First Piola-Kirchhoff stress tensor given in this section, the shear stress can be characterized with the following equation,

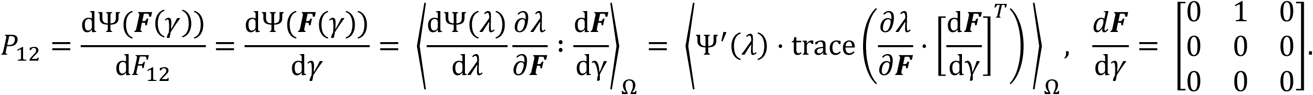

The fitting of the model’s parameters was carried out by means of the nonlinear least squares MATLAB function *lsqnonlin*. **Extended Data Figure 4**e shows a simple shear curve generated by this model in comparison to the experimental shear rheology curve (average of the one shown in **Figure 1**d) employed in the fitting of the corresponding material parameters up to a shear strain equivalent to 30%. The values obtained were: *κ*_*o*_= 167.2155 Pa, *d*_*o*_=0.375, *λ*_*s*_=0.0719, and *d*_s_=0.0180.

### Comparing DD-TFM with continuum

In principle, one could generate in silico displacement fields and forces (as done above to compare *HPBIM* with the *Classic* inverse method (**In silico ground truth simulations and validation**), to compare continuum TFM with DD-TFM. However, such simulations would require defining the ground truth using either a continuum finite-element model or a discrete fiber-network model, thereby inherently favoring one of the two methods. We therefore did not conduct a quantitative ground-truth comparison. Our goal was to compare the mechanical readouts that the two approaches derive from the same experimental displacement measurements: cell ranking based on force-related metrics, stress propagation away from the cell surface, and protrusion-level displacement decay.

#### Cell ranking differences (shown in **Figure 2f,g**)

To compare whether continuum TFM and DD-TFM ranked cells similarly according to their mechanical activity, we analyzed the subset of cells for which both methods were available. For each cell, three scalar metrics were computed: total force, strain energy and maximum force.

For continuum TFM, tractions and displacements were evaluated on the cell surface,*S*. Surface strain energy was computed as: 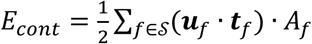, where *S*is the set of cell-surface faces, and ***u*** and ***t*** and *A*_*f*_ are the face-centered displacement, traction and area, respectively. Total force was computed as: 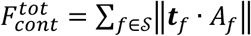, and maximum force was defined as: 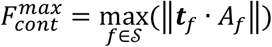.

For DD-TFM, nodal displacements and forces were summed over all *HPBIM* deformation increments. Metrics were then computed over the nodes closest to the cell surface, *C*. Strain energy and total force were defined as described in **Force recovery methods for DD-TFM**, and maximum force was defined as: 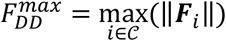.

For each metric, cells were ranked independently for continuum TFM and DD-TFM in descending order, so that rank 1 corresponded to the highest value. Rank agreement was visualized by plotting the continuum rank against the DD-TFM rank (**Figure 2**g); points on the diagonal indicate identical rankings between methods, whereas deviations from the diagonal indicate disagreement (color-coded). Spearman’s rank correlation was used to assess whether continuum TFM and DD-TFM ordered cells similarly (reported in **Figure 2**g). We also computed Pearson’s correlation to assess the linear association between the corresponding metric magnitudes, obtaining 0.53, 0.72 and 0.05 for total force, strain energy and maximum force, respectively.

#### Stress spatial decay (shown in **Extended Data Figure 4f**)

To compare the spatial decay of matrix stresses between continuum TFM and DD-TFM, we computed stress profiles as a function of distance to the cell surface for the subset of cells analyzed with both methods (**Extended Data Figure 4**f). For continuum TFM, we used the largest absolute principal stress at gel mesh nodes and computed the Euclidean distance from each gel node to the nearest cell-surface node. For DD-TFM, we used the total stress of each fiber element and computed the Euclidean distance from each element centroid to the nearest vertex of the segmented cell surface. Stress values were binned from 0 to 60 µm from the cell surface using 0.25 µm-wide bins and averaged. Profiles were then smoothed with a 5 µm moving average window. To compare the shape of stress decay independently of absolute stress magnitude, each profile was baseline-corrected by subtracting the median far-field stress between 50 and 60 µm from the cell surface. Negative values after baseline subtraction were set to zero. Each profile was then normalized by its maximum stress within the near-cell region, defined as distances ≤5 µm. The normalized DD-TFM and continuum profiles were plotted for individual cells and as mean and shaded standard deviation across cells (**Extended Data Figure 4**f).

The half-decay distance was defined as the distance from the cell surface at which the normalized stress profile first decreased below 0.5. When the threshold was crossed between two distance bins, the half-decay distance was estimated by linear interpolation between the adjacent bins. Because each cell was analyzed with both continuum TFM and DD-TFM, paired method differences were assessed using a two-sided Wilcoxon signed-rank test (MATLAB’s `signrank` function), which does not assume normally distributed paired differences.

#### Displacement decay from protrusions (shown in **Figure 2h,i** and **Extended Data Figure 4**g,h)

To compare displacement decay away from cell protrusions, we analyzed a representative cell for which measured displacement fields, continuum TFM, and DD-TFM displacement fields were available. We first segmented the cell protrusions using the approach described in **In silico ground truth simulations and validation** but using a cell-surface mesh instead of the cell binary image. Surface vertices at >10 µm from the inscribed sphere were classified as protrusion vertices. Protrusion vertices were grouped into connected clusters using the triangular surface connectivity; clusters containing fewer than 10 surface vertices were excluded. For each protrusion cluster, the protrusion direction was defined by its first principal component and oriented outward from the cell. The protrusion tip was then defined as the cluster vertex with the largest outward projected distance along this axis. Points in the hydrogel domain ahead of each protrusion were selected using a finite conical region originating at the protrusion tip. For a candidate hydrogel point ***x***, the vector from the protrusion tip ***c*** was computed as ***r*** = ***x*** − ***c***. Its axial projection along the protrusion unitary direction ***v*** was *s* = ***r*** · ***v*** and its radial distance from the cone axis (which coincides with the protrusion direction) was *ρ* = ‖***x*** − (***c*** + *s****v***)‖. The cone radius increased with axial distance according to *R*(*s*) = min(*R*_0_ + *s* tan *θ, R*_*max*_), with *R*_0_=10 µm, *θ*=30°, maximum axial length *L*_*max*_=30 µm, and maximum radius *R*_*max*_=30 µm. Candidate points were selected if they were located ahead of the protrusion and inside the cone: 0 < *s* ≤ *L*_*max*_ and *ρ* < *R*(*s*). In the measured and continuum TFM analysis, this criterion was applied to gel mesh nodes. In DD-TFM, the same criterion was applied to fiber-element centroids, after which both nodes belonging to the selected elements were included in the displacement-decay analysis.

For each selected point, the Euclidean distance to the protrusion tip was computed and used as the coordinate for the decay analysis (**Figure 2**i). Displacement magnitudes were extracted for the continuum TFM solution, the measured displacement field represented on the continuum mesh, and the DD-TFM solution. Each displacement field (for a certain protrusion) was normalized independently by the 95th percentile of displacement magnitudes within the proximal 0–10 µm region from the protrusion tip. Normalized displacement magnitudes were plotted as individual points against distance from the protrusion tip. To test how well each displacement field could be described by a power-law dependence on distance from the protrusion tip, normalized displacement magnitudes between 1 and 45 µm from the protrusion tip were fitted using shifted power-law models.

First, we fitted a shifted 1/r decay model: 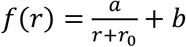. Here, *r* is the Euclidean distance to the protrusion tip, *a* is the fitted amplitude, *r*_0_ is a non-negative distance shift that avoids singular behavior near the protrusion tip, and *b* is a non-negative offset accounting for residual far-field displacement. Although the continuum modeling, we employed in this work is more complex, a 1/r decay in displacements is the profile of the displacement field a solid undergoes at a distance r from the application of a point force in a linear and elastic body (referred as Green’s solution). Fits were implemented in MATLAB using custom fitting functions based on `lsqcurvefit`. Fit quality was quantified using R^2^ and RMSE. To reduce sensitivity to local minima, fits were repeated from multiple initial guesses for *r*_0_ and, for the free-exponent model, for *n*. The solution with the lowest sum of squared residuals was retained. To ensure that the poorer fit of DD-TFM was not simply due to the larger number of sampled fiber nodes, we repeated the analysis over 1000 bootstrap iterations in which the DD-TFM point cloud was randomly downsampled to match the number of continuum/measured points before refitting the shifted 1/*r* model. The R^2^ and RMSE distribution values are reported in **Extended Data Figure 4**g, showing median values of R^2^ (~0.4) and RMSE (~0.1), close to the ones reported for the original DD-TFM point cloud (**Figure 2**i).

We also fitted a free-exponent shifted power-law model: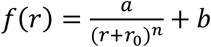. Because the fitted model contains a distance shift *r*_0_, the fitted exponent corresponds to decay with respect to *r* + *r*_0_, not directly with respect to the physical distance *r*. We therefore additionally reported an effective exponent, *n*_*eff*_, defined as the exponent of an equivalent unshifted power law over the plotted distance range *r*_1_=0.5 µm to *r*_2_=40 µm: 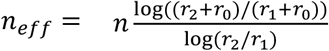. Interestingly, *n*_*eff*_~1 for the continuum, consistent with a decay that is approximately inverse to the distance. While both measured and DD-TFM fits yielded *n*_*eff*_<1, the goodness of fit was low (R^2^~0.3), indicating that these displacement fields are poorly described by a 1/r decay (**Extended Data Figure 4**h).

**Extended Data Figure 4:**
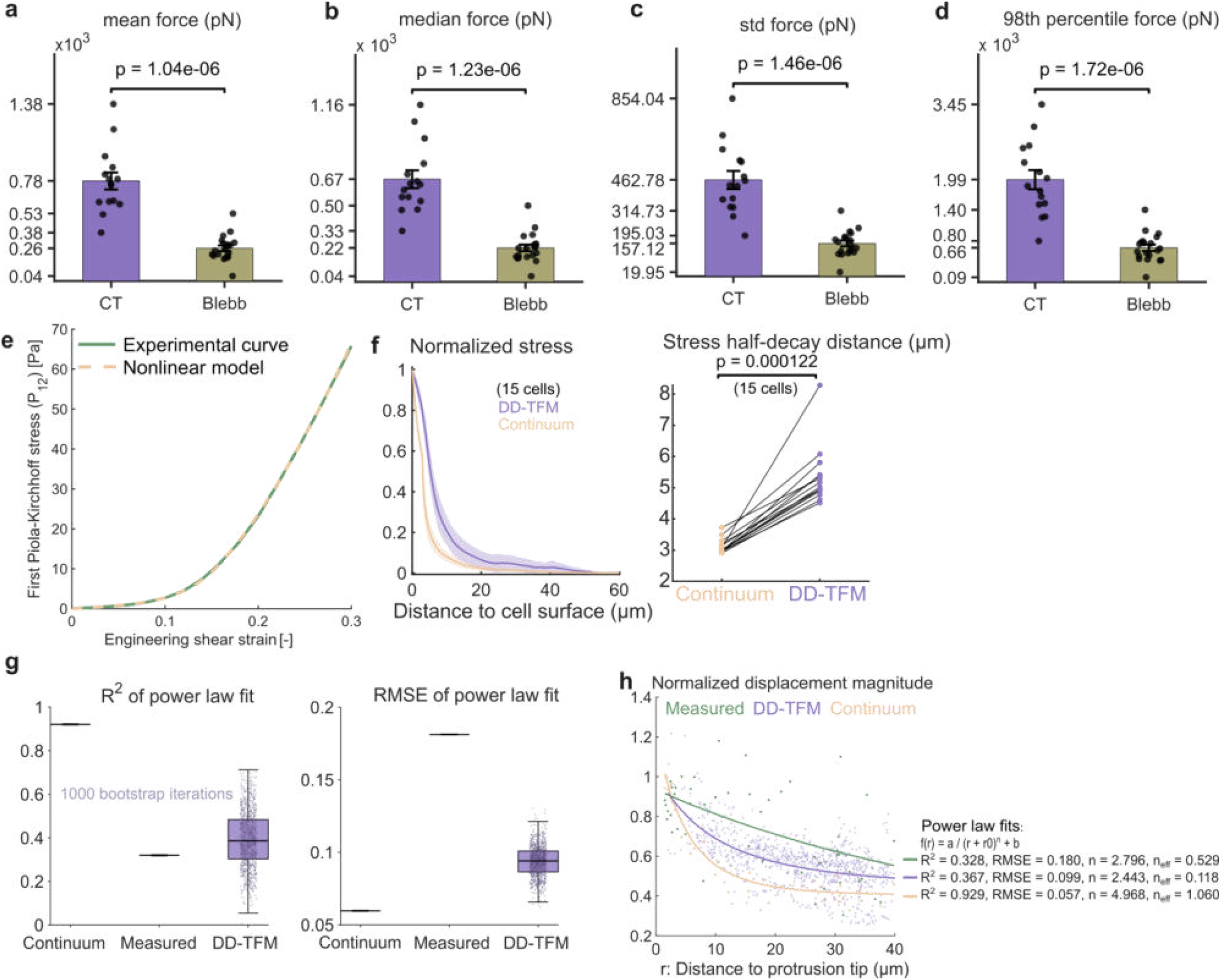
Additional experimental DD-TFM readouts and comparison with continuum TFM. **a–d:** Additional force-magnitude metrics recovered (in addition to **Figure 2**b) with DD-TFM in control (CT) and blebbistatin-treated (Blebb) HUVECs. Metrics include mean, median, standard deviation and 98th percentile of force magnitude at the fiber nodes closest to the cell surface. Bars show mean ± SEM, with individual cells overlaid as scatter points. Statistical comparisons were performed using a two-sided Wilcoxon rank-sum test. **e:** Fitting of experimental curve of 1.5 mg/ml collagen hydrogel with the proposed nonlinear hyperelastic model for shear rheology tests. The experimental curve results from the average of the 3 samples shown in **Figure 1**d. **f:** Comparison of normalized stress-decay profiles between DD-TFM (purple) and continuum (orange) TFM for the subset of 15 cells analyzed with both methods (left) and corresponding half-decay distance comparison (right). Lines and shading indicate mean ± SD. Statistics used a two-sided Wilcoxon signed-rank test. **g:** Matched-sampling analysis of the protrusion displacement-decay fit. The DD-TFM point cloud was repeatedly downsampled to match the number of continuum/measured points, and the shifted 1/r model was refitted over 1000 iterations. Distributions of R^2^ (left) and RMSE (right) show that the poorer DD-TFM fit was not explained by the larger number of sampled fiber nodes. **h:** Free-exponent shifted power-law fits of normalized displacement magnitude as a function of Euclidean distance from the protrusion tip for measured (green), continuum TFM (orange) and DD-TFM (purple) displacement fields. Effective exponents, n_eff_, were computed over the plotted distance range to account for the fitted distance shift.

## Acknowledgements

Colormaps *batlow, roma, lajolla* and *acton* from the Scientific Colourmaps toolbox^53^ were used throughout this manuscript. J.B-F. gratefully acknowledges the support from Research Foundation Flanders (FWO) junior postdoctoral fellowship (1259223N). G.H.K. acknowledges funding from the Convergence programme Syn-Cells for Health(care) under the theme of Health and Technology. J.A.S.H. acknowledges project MCIU/AEI/10.13039/501100011033 [PID2024-161310NB-I00]. H.V.O acknowledges KU Leuven grants iBOF/21/083C and C14/24/112, FWO project G0C2422N and ERANET NEURON JTC 2022 call – project G0L1522N.

## Author contributions

J.B.F., J.A.S.H. and H.V.O conceptualized the study. J.B.F. conducted the study and wrote the manuscript. J.B.F. and L.K. performed TFM experiments. J.B.F., A.A.F., E.N.O. and J.A.S.H developed all computational methods. A.A.F. computed TFM results using the continuum model. J.B.F., L.K. and D.L. performed shear rheology with guidance from I.M., G.K., and R.C. N.V, A.S. and H.V.O assisted with computational and experimental design. All authors reviewed and gave feedback on the manuscript.

